# Cysteine induces mitochondrial reductive stress in glioblastoma through hydrogen peroxide production

**DOI:** 10.1101/2021.09.10.459864

**Authors:** Evan K Noch, Laura Palma, Isaiah Yim, Daniel Barnett, Alexander Walsh, Bhavneet Bhinder, Elisa Benedetti, Jan Krumsiek, Justin Gurvitch, Sumaiyah Khwaja, Olivier Elemento, Lewis C. Cantley

**Affiliations:** Department of Neurology, Division of Neuro-oncology, New York, NY, USA; Sandra and Edward Meyer Cancer Center, Weill Department of Medicine, New York, NY, USA; Graduate School of Medical Sciences, New York, NY, USA; Department of Physiology and Biophysics, Institute for Computational Biomedicine, New York, NY, USA; Caryl and Israel Englander Institute for Precision Medicine, Weill Cornell Medicine, New York, NY, USA

**Keywords:** Cysteine, N-acetylcysteine, cysteine susceptibility, glioma, glioblastoma, metabolism, reductive stress, hydrogen peroxide, glucose dependence

## Abstract

Glucose and amino acid metabolism are critical for glioblastoma (GBM) growth, but little is known about the specific metabolic alterations in GBM that are targetable with FDA-approved compounds. To investigate tumor metabolism signatures unique to GBM, we interrogated The Cancer Genome Atlas for alterations in glucose and amino acid signatures in GBM relative to other human cancers and found that GBM exhibits the highest levels of cysteine and methionine pathway gene expression of 32 human cancers. Treatment of patient-derived GBM cells with the FDA-approved cysteine compound N-acetylcysteine (NAC) reduce GBM cell growth and mitochondrial oxygen consumption, which was worsened by glucose starvation. Mechanistic experiments revealed that cysteine compounds induce rapid mitochondrial H_2_O_2_ production and reductive stress in GBM cells, an effect blocked by oxidized glutathione, thioredoxin, and redox enzyme overexpression. These findings indicate that GBM is uniquely susceptible to NAC-driven reductive stress and could synergize with glucose-lowering treatments for GBM.

## Introduction

Glioblastoma (GBM) remains one of the most lethal human cancers, with a median overall survival of just 14-16 months despite surgical resection, chemotherapy, and radiation (Stupp et al., 2005; Stupp et al., 2015). Increased flux through glucose and amino acid metabolic pathways is central to GBM proliferation (Zhu and Thompson, 2019) and is regulated primarily through oncogene-controlled membrane transporters and metabolic enzymes (Barthel et al., 1999; Ilic et al., 2017; Yang et al., 2012). These nutrients facilitate energy production through ATP generation and also protect against oxidative stress through the generation of NADPH (Zhu and Thompson, 2019).

Unlike systemic cancers, GBM is adapted to the nutrient pool available within the central nervous system. Because the brain uses 20% of the body’s glucose (Mink et al., 1981), GBM cells can access a vast supply of glucose for growth. In these cells, glucose is preferentially used for mitochondrial glucose oxidation, indicating a dependency on mitochondrial-generated ATP production in GBM (Marin-Valencia et al., 2012). The excess of certain amino acids, such as glutamate, in the brain also fuels GBM growth by killing surrounding neurons, promoting inflammation, and inducing brain edema (Long et al., 2019; Noch and Khalili, 2009; Venkataramani et al., 2019; Venkatesh et al., 2019; Ye and Sontheimer, 1999). These studies support the targeting of brain-associated nutrient metabolism in GBM.

Cysteine is a sulfur-containing amino acid that is required for glutathione production. Cysteine levels are regulated through a variety of membrane-bound amino acid transporters, including the cystine-glutamate antiporter system Xc^−^, which plays a critical role in the oxidative stress response in GBM and other cancers (Chung et al., 2005; Robert et al., 2015). The oxidation and reduction status of intracellular cysteine is also an indicator of the cellular redox status and can control survival from oxidative stress (Banjac et al., 2008). Likewise, the intracellular redox environment can greatly affect the function of metabolic enzymes through direct interaction with reactive cysteine residues (Fu et al., 2017; Kuljanin et al., 2021; Weerapana et al., 2010). Yet, the role of cysteine in mediating nutrient metabolism and cell growth in GBM remains largely unexplored.

We demonstrate that GBM exhibits a unique susceptibility to the amino acid, cysteine, and a variety of compounds that contain cysteine. This susceptibility is the result of direct mitochondrial toxicity, produced by paradoxical reductive stress that triggers production of mitochondrial hydrogen peroxide. This toxicity is exacerbated by glucose deprivation, making strategies that lower serum glucose, including anti-hyperglycemic agents and the ketogenic diet, potentially synergistic with cysteine-containing compounds to treat GBM

## Results

### L-cysteine induces cytotoxicity in patient-derived glioma cells

As a precursor to the antioxidant glutathione, cysteine protects against harmful reactive oxygen species (ROS) that induce oxidative stress, inflammation, and aging (Bavarsad Shahripour et al., 2014). Likewise, in cancer, cysteine can fuel tumor growth (Combs and DeNicola, 2019). However, recent studies have demonstrated that cysteine compounds induce hydrogen sulfide production, reductive mitochondrial stress, and apoptosis (Finn and Kemp, 2012; Kolossov et al., 2015; Tsai et al., 1996; Yang et al., 2013; Zhang et al., 2012). These findings indicate a context-specific role of cysteine-promoting compounds on cell survival in cancer.

We examined the cysteine and methionine pathway in 2 patient-derived glioma cell lines (603, anaplastic oligodendroglioma; and 667, GBM, IDH wild-type). 603 cells contain 1p/19q co-deletion and IDH1 R132 mutation, and 667 cells contain Chromosome 7 gain, *pten* deletion, *cdkn2a/2b* loss, and *pdgfra* amplification. These cells are grown in the absence of serum but with growth factors (EGF and FGF) to better recapitulate the tumor environment. We starved these cells of cysteine and methionine and added increasing doses of L-cysteine and L-methionine to the tissue culture medium. We found that these cells grew less when starved of cysteine and methionine and paradoxically, also when treated with > 5 mM L-cysteine (Figure 1A, C). Addition of L-methionine did not have a major effect on growth (Figure 1B). Treatment with L-cysteine and L-methionine in combination recapitulated the effects of L-cysteine treatment alone.

**Figure 1.**
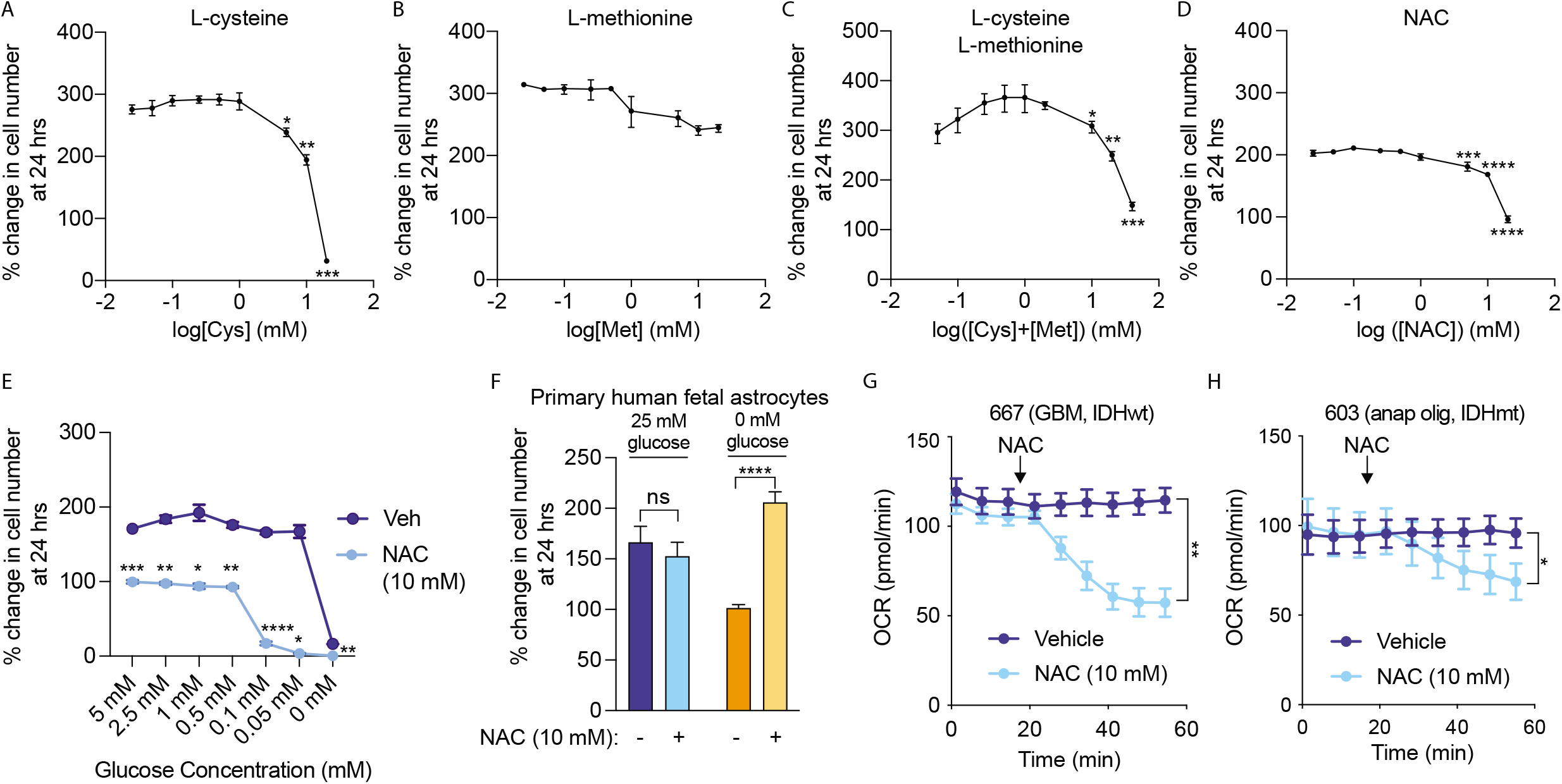
Cysteine induces cytotoxicity and reduces oxygen consumption in glioma cells. 667 cells were starved of cysteine and methionine, and growth was measured after treatment with L-cysteine (A), L-methionine (B), or the combination (C). 667 cells in cysteine- and methionine containing media were treated with 10 mM NAC, and growth was measured at 24 hours (D). 667 cells were treated with vehicle or 10 mM NAC and subjected to the indicated glucose concentrations (0-5 mM) for growth measurement at 24 hours (E). Primary human fetal astrocytes were treated with 10 mM NAC under normal glucose conditions (25 mM) or glucose starvation (0 mM), and growth was measured at 24 hours (F). 603 cells (G) and 667 cells (H) were treated with 10 mM NAC, and oxygen consumption rate (OCR) was measured using the Agilent Seahorse assay. *, p < 0.05, **, p < 0.01, ***, p < 0.001, ****, p < 0.0001.

N-acetylcysteine (NAC), an FDA-approved drug to treat acetaminophen overdose and asthma, is a cysteine-containing prodrug that possesses greater solubility than L-cysteine under physiological conditions. We chose to use NAC in subsequent experiments as a putative cysteine-containing compound. NAC administration resulted in reduced growth at doses of 5, 10, and 20 mM (Figure 1D). L-cysteine, L-cystine, and D-cysteine also reduced growth of 667 cells, as did reduced glutathione (GSH) and glutathione reduced ethyl ester (GREE), which more readily crosses the cell membrane (Figure S1A-E). Inclusion of serum in growth media did not affect NAC-induced growth inhibition (Figure S1F). NAC treatment also exhibited a dose-dependent toxicity with glucose starvation, with a marked reduction in growth noted at glucose concentrations less than 0.5 mM (Figure 1E). 2-deoxyglucose (2-DG), which prevents glucose oxidation and subsequent entry into glycolysis, recapitulated the synergistic effect of glucose starvation on NAC-induced cytotoxicity (Figure S1G). Notably, primary human fetal astrocytes (PHFA) were unaffected by NAC treatment (Figure 1F). In fact, NAC treatment improved growth when PHFA were starved of glucose. Likewise, non-glioma cells, including A549 (lung), H1975 (lung), HCC2935 (lung), MCF-7 (breast), and HT-29 (colon) cells were all unaffected by NAC treatment (Figure S1H), demonstrating a unique susceptibility of GBM to cysteine compounds.

Since cysteine compounds have been shown to modulate mitochondrial metabolism in cancer cells (Kolossov et al., 2015), we examined mitochondrial oxygen consumption in the presence of NAC. Treatment with 10 mM NAC induced rapid reduction in mitochondrial oxygen consumption in 667 and 603 cells (Figure 1G-H). L-cysteine, D-cysteine, GSH, and GREE also caused a similar reduction in OCR; the effect of GREE was likely more pronounced than GSH because of more efficient cell entry (Figure S1I-L). A549, H1975, and HCC2935 cells exhibited no reduction in OCR (Figure S2A-C).

### Cysteine and methionine metabolic genes are upregulated in glioblastoma

To determine whether GBM exhibits unique cysteine metabolism as compared to other human cancers, we conducted a Kyoto Encyclopedia of Genes and Genomes (KEGG) analysis on glucose and amino acid pathways in GBM using The Cancer Genome Atlas (TCGA) database. We found that GBM exhibits the highest expression of cysteine and methionine pathway gene expression of 32 human cancers and that this gene expression is increased in GBM versus WHO Grade II and III glioma (Figure 2A-B, Figure S3A). There was no significant difference in the cysteine-methionine signature among brain regions (Figure S3B), and normal brain tissue exhibited a similar level of this signature as other organs, with liver having the highest level (Figure S3C).

**Figure 2.**
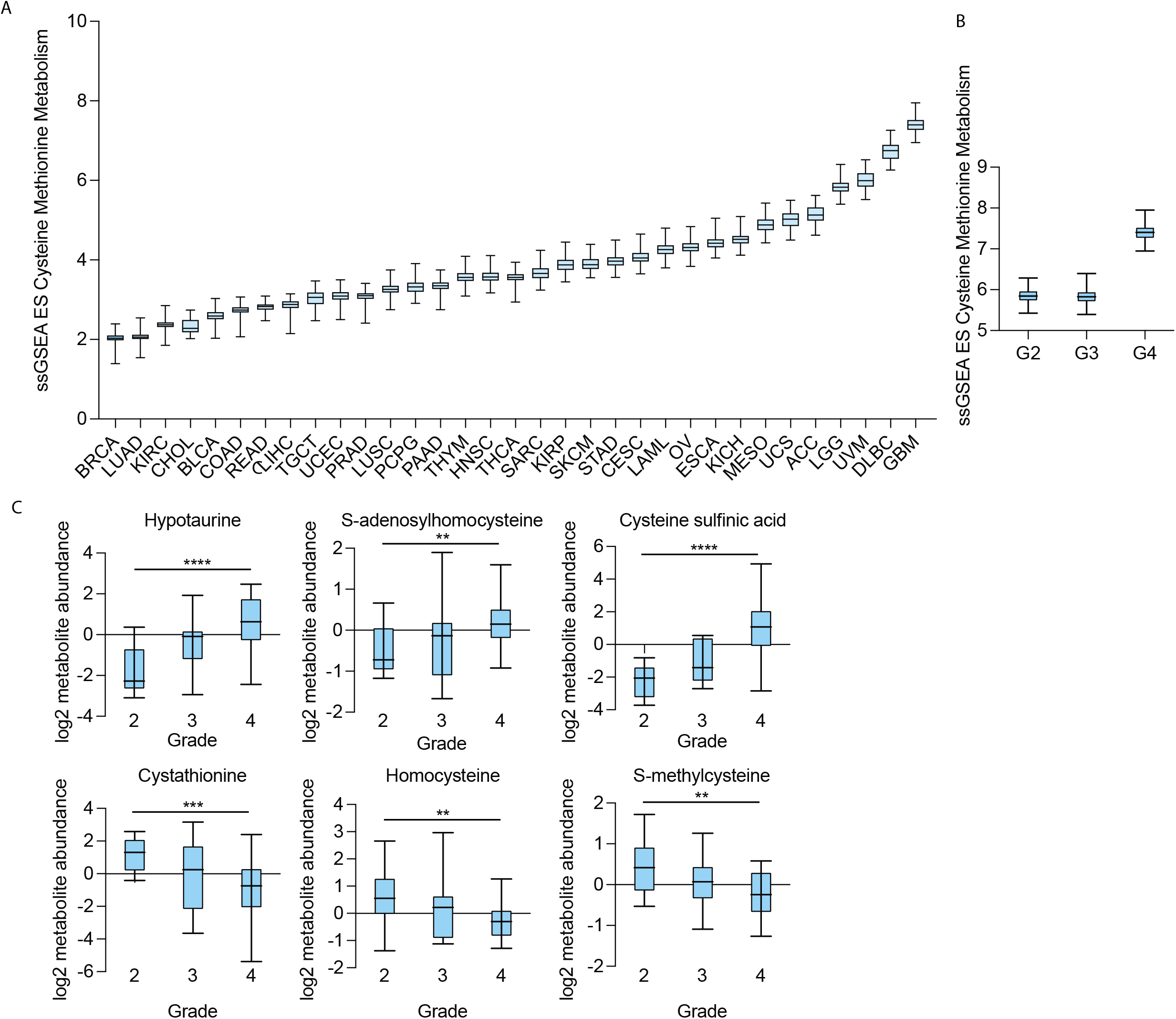
Cysteine and methionine genes and metabolites are overexpressed in glioma. (A) Cysteine-methionine KEGG signature in 32 human cancers obtained from The Cancer Genome Atlas (TCGA). (B) Cysteine-methionine KEGG signature in glioma based on WHO Grade. (C) Cysteine metabolite abundance was obtained from the cancer metabolite database (Chinnaiyan et al., 2012), and significantly different metabolites among WHO Grade 2, 3, and 4 gliomas are presented. **, p < 0.01, ***, p < 0.001, ****, p < 0.0001.

We next examined metabolite levels in human glioma using the Pan-cancer Metabolism Data Explorer. When we compared metabolite levels among WHO Grade 2, 3, and 4 gliomas, we found that hypotaurine, S-adenosylhomocysteine, and cysteine sulfinic acid increased with grade, whereas cystathionine, homocysteine, and S-methylcysteine decreased with grade (Figure 2C). These findings reinforce that cysteine metabolic genes are not only upregulated in GBM but also contribute to alterations in levels of cysteine metabolites.

### N-acetylcysteine induces mitochondrial toxicity in glioma cells

Because NAC caused a rapid reduction in mitochondrial oxygen consumption in GBM cells, we characterized additional metabolic deficits induced by NAC. NAC caused a reduction in mitochondrial membrane potential within 6 hours of treatment (Figure 3A). Likewise, NAC induced a reduction in the NADPH/NADP+ ratio and reduced glutathione (GSH)/oxidized glutathione (GSSG) ratio within 5 minutes of treatment. Even though NAC is a known antioxidant, GBM cells exhibited a rapid oxidized state with NAC treatment, indicating a differential response in these cells. We confirmed this finding by treating 667 cells with 10 mM ^13^C3-L-cysteine for 5 minutes and measuring metabolic flux. We found that ^13^C3-L-cysteine treatment predominantly led to labeled L-cystine, L-cystathionine, and reduced and oxidized glutathione (Figure S4A). Interestingly, we found high levels of oxidized glutathione and low levels of reduced glutathione in treated cells, suggesting that cysteine treatment led to a rapid reduction in the GSH/GSSG ratio. Metabolic flux analysis using U-^13^C6-glucose (10 mM) in the presence of 10 mM NAC for 5 minutes demonstrated that upper glycolysis was not largely affected by NAC (Figure S4B). In fact, lower glycolytic metabolites, such as pyruvate (m+3) and lactate (m+3), and 3-phosphoglycerate derivates, such as serine (m+3) and glycine (m+2) were significantly reduced, suggesting that glucose may be shunted towards the pentose phosphate pathway to produce NADPH that would facilitate glutathione production.

**Figure 3.**
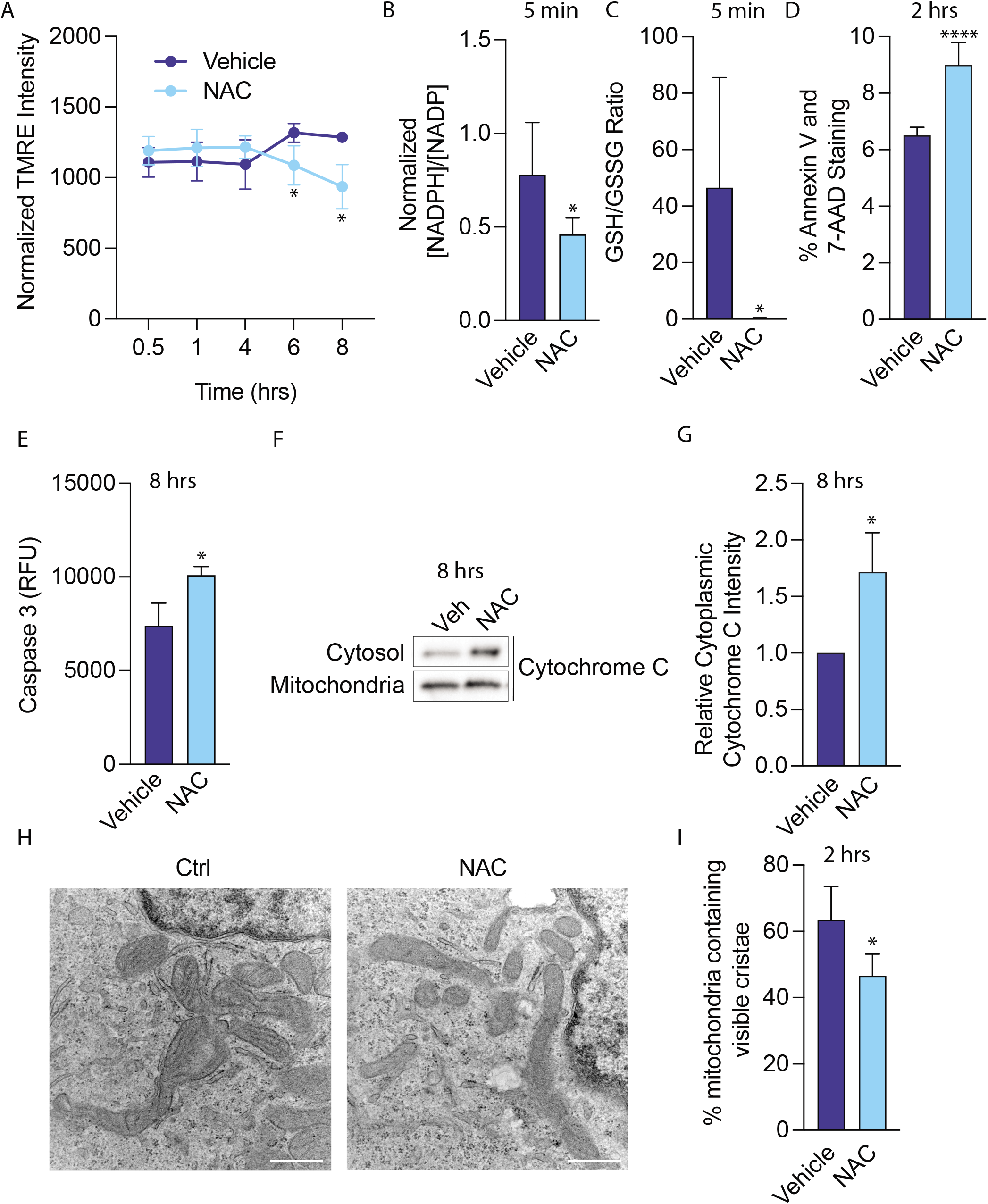
NAC induces mitochondrial toxicity in glioma cells. (A) 667 cells were treated with 10 mM NAC, and mitochondrial membrane potential was measured using tetramethylrhodamine (TMRE). 667 cells were treated with 10 mM NAC for 5 minutes, and NADPH/NADP+ levels (B) and reduced glutathione (GSH)/oxidized glutathione (GSSG) (C) was measured. (D) 667 cells were treated with 10 mM NAC for 2 hours, and Annexin V and 7-AAD levels were measured by flow cytometry. (E) 667 cells were treated with 10 mM NAC for 8 hours, and activated Caspase 3 levels were measured by fluorescence. (F) 667 cells were treated with 10 mM NAC for 8 hours, and cytochrome C release from the mitochondria to the cytosol was measured through mitochondrial immunoprecipitation and subsequent mitochondrial and cytoplasmic lysate preparation followed by immunoblot. Quantification of 3 independent experiments is shown in (G). (H) 667 cells were treated with 10 mM NAC for 2 hours, and cells were prepared for transmission electron microscopy. The number of visible cristae per mitochondria were counted from 312.5 μm^2^ area and quantified in (I). Results are indicative of 4 biological replicates. Scale bars = 500 nm. *, p < 0.05, ****, p < 0.0001.

Cells treated with NAC exhibited significantly higher levels of Annexin V and 7-aminoactinomycin D (7-AAD), indicators of apoptosis and necrosis, respectively (Figure 3D). 8-hour NAC treatment also increased activated caspase 3 levels (Figure 3E) and induced cytochrome c released into the cytosol, an indication of mitochondrial membrane damage (Figure 3F-G). Finally, 2-hr NAC treatment led to dissolution of mitochondrial cristae as assessed by transmission electron microscopy (Figure 3H-I).

### NAC induces hydrogen peroxide and reductive stress in glioma cells

Prior studies have shown that reducing agents like NAC induce reductive stress by depleting oxidized electron acceptors (glutathione and thioredoxin), thereby allowing reduction of O_2_ to H_2_O_2_ (Korge et al., 2015). Given the rapid reduction in OCR induced by NAC and given the sensitivity of mitochondrial respiration to changes in redox state, we hypothesized that the mechanism of NAC toxicity involved a rapid change in the mitochondrial redox state that induces mitochondrial toxicity. We isolated GBM mitochondria from 667 cells using a well-validated HA-tag-based method (Chen et al., 2017) and demonstrated immunoprecipitation of mitochondria with the matrix marker, citrate synthase (Figure S5A-B). Using the fluorescent probe, Amplex UltraRed, we measured H_2_O_2_ production in isolated alamethicin-permeabilized mitochondria and found that NAC rapidly increased the rate of H_2_O_2_ production, which was suppressed by H_2_O_2_ detoxification with catalase (Figure 4A). Though antioxidants like NAC are known to induce spontaneous H_2_O_2_ production in tissue culture medium and buffers, rates of H_2_O_2_ production were higher in NAC-treated cells than in mitochondrial assay buffer alone (Figure S6A). In addition, NAC induced a similar reduction in OCR as 100 μM H_2_O_2_ (Figure S6B), indicating that NAC produces higher levels of H_2_O_2_ than have been shown to be produced in tissue culture medium alone. We also measured mitochondrial H_2_O_2_ production by direct visualization using the Mito-PY1 probe, a mitochondrial permeable reactive oxygen species fluorescent probe that specifically reports mitochondrial H_2_O_2_. 2-hour treatment with NAC led to significantly higher levels of mitochondrial H_2_O_2_ than vehicle (Figure 4B-C). Together, these results indicate that NAC induces rapid mitochondrial H_2_O_2_ production in GBM cells, leading to eventual cytotoxicity.

**Figure 4.**
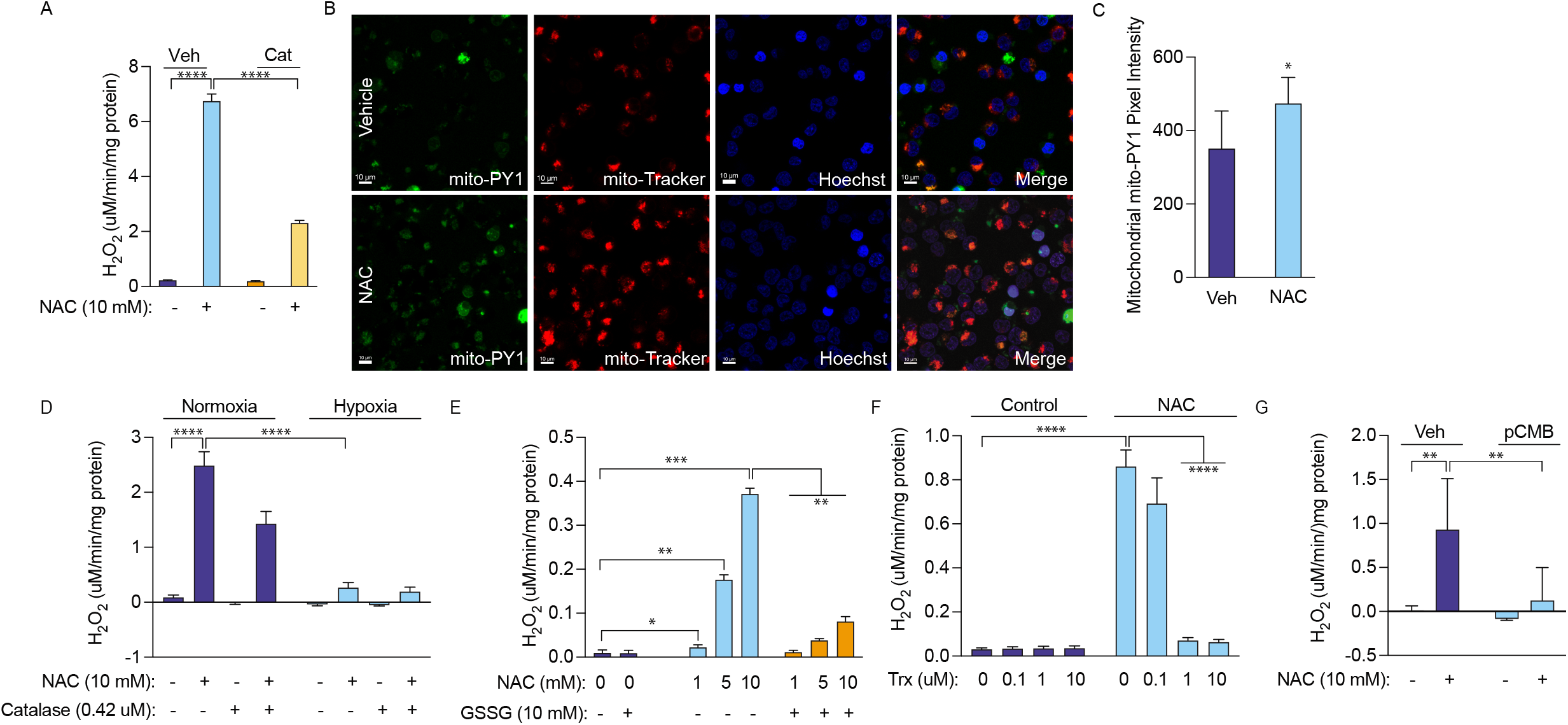
NAC induces hydrogen peroxide production in glioma cells. (A) Mitochondria were isolated from 667 cells, permeabilized with 10 μM alamethicin, and treated with vehicle or 10 mM NAC. The rate of H_2_O_2_ production was measured, and 0.42 μM catalase was used to detoxify H_2_O_2_. (B) 667 cells were treated with 10 mM NAC for 2 hours, and mitochondrial H_2_O_2_ was measured by confocal microscopy using the mito-PY1 dye. Mitotracker was used to determine mitochondrial localization. Mito-PY1 intensity was quantified in (C), with results indicating 3 independent experiments. 667 mitochondria were treated as in (A) but exposed to normoxia or 1% hypoxia in a modular hypoxia chamber (D), treated with oxidized glutathione (GSSG) (E), recombinant human Trx (F), or 50 μM perchlorobenzoic acid (pCMB) (G), after which time the rate of H_2_O_2_ production was measured. *, p < 0.05, **, p < 0.01, ***, p < 0.001, ****, p < 0.0001.

Because oxygen serves as an electron acceptor to facilitate reductive stress, we examined the effect of hypoxia (1% O_2_) on NAC-driven H_2_O_2_ production. When we treated isolated mitochondria with NAC and exposed them to normoxia or immediate hypoxia, we found significantly reduced H_2_O_2_ production (Figure 4D). These findings indicate that NAC-induced mitochondrial reductive stress likely occurs through electron overflow that converts oxygen into an alternate electron acceptor.

Electron acceptor deficiency is a critical determinant of reductive stress (Korge et al., 2015). By providing alamethicin-permeabilized mitochondria from 667 cells with oxidized glutathione (GSSG) or recombinant thioredoxin (Trx), we nearly completely rescued NAC-induced H_2_O_2_ production (Figure 4E-F). Thiol-reactive compounds can also alter electron flow from reducing agents to available acceptors. Because we hypothesize that NAC causes reduction of reactive thiols, co-administration of thiol-reactive agents should decrease the effect of NAC. We found that p-chloromercuribenzoic acid (pCMB) decreased NAC-induced H_2_O_2_ production (Figure 4G). Thus, a shortage of oxidized electron acceptors mediates cysteine susceptibility in GBM cells.

### Redox enzyme overexpression rescues NAC-induced mitochondrial H_2_O_2_ production

The glutathione/glutathione peroxidase 3 (Gpx)/glutathione reductase (GR) and the peroxiredoxin 3 (Prx)/thioredoxin 2 (Trx)/thioredoxin reductase (TrxR2) systems are the major mitochondrial matrix H_2_O_2_ scavenging systems and are regulated by the redox environment within the matrix. We hypothesized that cysteine susceptibility in GBM is due to a specific deficiency in mitochondrial redox enzymes. Therefore, we investigated the effect of genetic overexpression of redox enzymes on NAC-induced cytotoxicity and H_2_O_2_ production. Thioredoxin-2 (Trx2) is the mitochondrial thioredoxin that acts as the electron acceptor for mitochondrial thioredoxin reductase (TrxR2). We genetically overexpressed cytoplasmic Trx and mitochondrial Trx2 in our GBM cells (Figure S7A-B) and found that Trx2 overexpression accelerated GBM growth in untreated cells and reduced cytotoxicity after NAC treatment (Figure 5A). Likewise, Trx2 overexpression rescued NAC-induced H_2_O_2_ production (Figure 5B).

**Figure 5.**
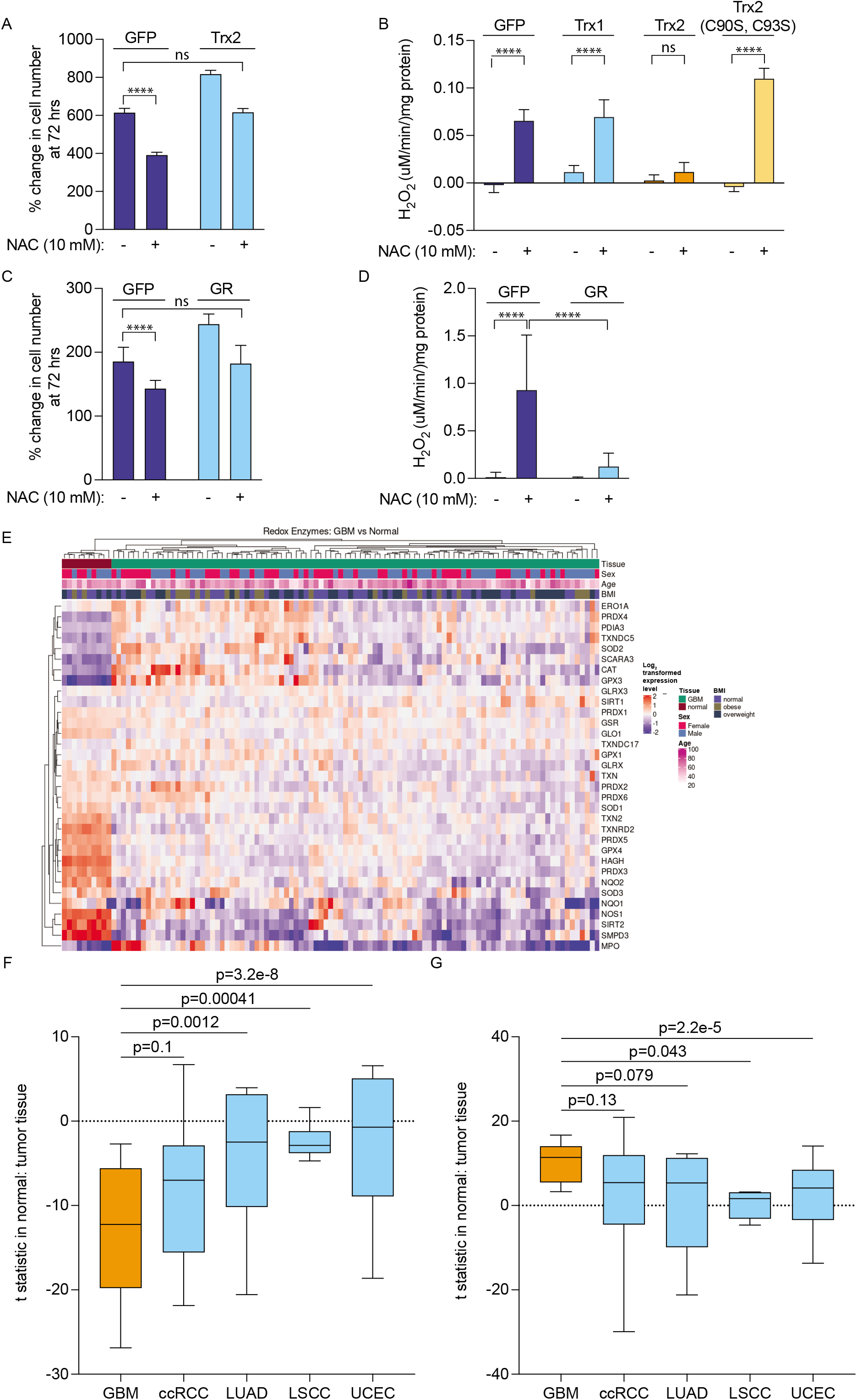
Redox enzyme overexpression ameliorates NAC-induced reductive stress in GBM. (A) GFP- and Trx-2-expressing 667 cells were treated with 10 mM NAC for 72 hours, and growth was measured. (B) Mitochondria were isolated from 667 cells expressing GFP, Trx1, Trx2, or Trx2 C90S/C93S, permeabilized with 10 μM alamethicin, and treated with vehicle or 10 mM NAC, after which time the rate of H_2_O_2_ production was measured. (C) GFP- and GR expressing 667 cells were treated with 10 mM NAC for 72 hours, and growth was measured. (D) Mitochondria were isolated from 667 cells expressing GFP or GR, permeabilized with 10 μM alamethicin, and treated with vehicle or 10 mM NAC, after which time the rate of H_2_O_2_ production was measured. (E) GBM and normal brain tissue proteomics data were obtained from CPTAC database, and the database was interrogated based on a signature of 39 known redox enzymes. An unsupervised hierarchically clustered heatmap was generated. A signature of either significantly downregulated (F) or significantly upregulated (G) proteins was assayed among GBM, clear cell renal cell carcinoma, lung squamous cell carcinoma, lung adenocarcinoma, and uterine corpus endometrial carcinoma available from CPTAC. ****, p < 0.0001.

Trx and Trx2 contain a dithiol active site motif (Cys-Gly-Pro-Cys) that is conserved from archaea to mammals. In Trx2, these cysteine residues occur at amino acid positions 90 and 93. To study the importance of these cysteine residues in the electron acceptor function of Trx2, we mutated these residues to serine, thereby abrogating oxidative reactivity. We found that mutation of both cysteine residues to serine abolished Trx2’s rescue of H_2_O_2_ production, indicating that the enzymatic function of Trx2 is critical for its rescue of reductive stress. Trx overexpression did not affect H_2_O_2_ production.

Glutathione reductase (GR) is an NADPH-dependent oxidoreductase enzyme containing FAD and a redox disulfide catalytic domain. GR regenerates GSH from GSSG to detoxify ROS but also facilitates reductive stress when GSSG is in short supply. Therefore, we sought to investigate the effect of GR overexpression on reductive stress in GBM cells. We genetically overexpressed GR (Figure S7C) and found that this overexpression accelerated GBM growth and reduced NAC-induced cytotoxicity. Similar to Trx2 overexpression, GR overexpression also nearly completely blocked NAC-induced H_2_O_2_ production. These findings indicate that a relative GR deficiency may promote reductive stress in response to NAC.

To determine whether GBM exhibits a relative deficiency of redox enzymes in comparison to other cancers, we investigated the National Cancer Institute’s Clinical Proteomic Tumor Analysis Consortium (CPTAC) using a redox enzyme signature composed of 39 proteins known to be involved in redox enzyme function. From this analysis, we found that GBM exhibited a lower redox enzyme signature than normal brain (Figure 5E). Within this redox enzyme signature, 19 proteins were significantly downregulated and 10 were significantly upregulated compared to normal brain tissue (Table 1). txn2 (Trx2), txnrd2 (TrxR2), and gsr (GR) were all significantly downregulated in GBM tissue as compared to normal brain.

**Table 1**
| **Gene Name** | **P-value** | **Median Difference in Expression** | **Direction of Regulation** |
| --- | --- | --- | --- |
| CAT | 1.01E-05 | 0.85 | up |
| GPX3 | 2.34E-07 | 1.70 | up |
| MPO | 0.0236873 | 0.91 | up |
| PDIA3 | 1.03E-06 | 0.61 | up |
| PRDX4 | 2.93E-07 | 1.00 | up |
| SCARA3 | 1.14E-06 | 1.15 | up |
| SIRT1 | 0.0003511 | 0.35 | up |
| SOD2 | 0.0019225 | 0.70 | up |
| TXNDC17 | 0.0011165 | 0.25 | up |
| TXNDC5 | 4.82E-07 | 0.83 | up |
| GLO1 | 2.62E-07 | -0.60 | down |
| GLRX | 0.0318177 | -0.30 | down |
| GPX4 | 2.21E-07 | -1.16 | down |
| GSR | 4.97E-05 | -0.31 | down |
| HAGH | 2.09E-07 | -1.62 | down |
| NOS1 | 3.27E-07 | -2.06 | down |
| NQO2 | 9.73E-07 | -1.12 | down |
| PRDX1 | 0.0060504 | -0.29 | down |
| PRDX2 | 0.0099638 | -0.45 | down |
| PRDX3 | 2.09E-07 | -1.39 | down |
| PRDX5 | 2.09E-07 | -1.06 | down |
| PRDX6 | 0.0066567 | -0.24 | down |
| SIRT2 | 8.29E-07 | -2.30 | down |
| SMPD3 | 2.48E-07 | -2.63 | down |
| SOD1 | 7.57E-06 | -0.63 | down |
| SOD3 | 0.0008276 | -0.74 | down |
| TXN | 6.21E-06 | -0.59 | down |
| TXN2 | 2.93E-07 | -0.86 | down |
| TXNRD2 | 2.09E-07 | -1.34 | down |

We then developed a signature containing either all significantly upregulated or all significantly downregulated proteins in GBM as compared to 4 other cancers whose proteomic data was available from CPTAC, including clear cell renal cell carcinoma (ccRCC), lung squamous cell carcinoma (LSCC), lung adenocarcinoma (LUAD), and uterine corpus endometrial carcinoma (UCEC). Among the downregulated protein signature, GBM’s signature was significantly lower than 3 of the other tumor types (p-value for ccRCC was 0.1). Among the upregulated protein signature, GBM’s signature was significantly higher than 2 of the other tumor types (p-value for LUAD and ccRCC was 0.079 and 0.13, respectively). These proteomics results suggest that deficiency of specific redox enzymes, including 3 of the most common enzymes involved in redox susceptibility, may underlie GBM’s unique susceptibility to cysteine.

## Discussion

Altered redox homeostasis is a hallmark of cancer. Cancer cells must maintain sufficient ROS to grow by inactivating phosphotyrosine phosphatases to facilitate receptor tyrosine kinase signaling but minimizing excessive ROS that hampers mitochondrial biogenesis. Though oxidative stress has been well explored in cancer, reductive stress is being increasingly recognized as an alternative mechanism of ROS generation that involves reductive overflow to alternate electron acceptors, such as O_2_ (Korge et al., 2015).

We found that the amino acid cysteine induces cytotoxic stress in GBM cells, driven by rapid reduction of oxygen consumption, depletion of intracellular NADPH and glutathione, and mitochondrial cristae dissolution. In GBM mitochondria, the cysteine prodrug NAC induced rapid H_2_O_2_ production, an effect blocked by catalase, hypoxia, oxidized glutathione, and thioredoxin. In other cancer cell lines, including breast, lung, colon, and pancreas, NAC had no effect on growth or oxygen consumption. Though NAC is mostly known as a cellular antioxidant, these results demonstrate that its function depends on the cell type and state in which it is used. For example, the process of reductive stress depends on excess mitochondrial reducing equivalents that overwhelm available electron acceptors. As a result, cells with sufficient electron acceptors will exhibit resistance to reductive stress. In our GBM cells, genetic overexpression of several redox enzymes, including Trx2 and GR, rescued NAC-induced H_2_O_2_ production, suggesting that redox enzyme deficiency supports cysteine susceptibility in GBM. Indeed, from an analysis of CPTAC data, we found that GBM exhibits lower levels of Trx2, TrxR2, and GR as compared to normal brain and other human cancers.

These results also add to a growing body of literature investigating cysteine metabolism in cancer. Most recent studies evaluating cysteine metabolism in cancer have focused on the dependency of cysteine for glutathione biosynthesis to buffer ROS, which is largely driven by the cystine-glutamate antiporter, system Xc (Briggs et al., 2016; Koppula et al., 2017; LeBoeuf et al., 2020; Lin et al., 2020; Shin et al., 2017). In this context, either extracellular cysteine depletion or intracellular glutamate depletion sensitizes cells to oxidative stress and subsequent death. Alternatively, some groups have found that cysteine abundance leads to cell death. For example, Prabhu et al. found that cysteine-containing compounds, such as cysteine sulfinic acid (CSA), cause cytotoxicity in GBM cells, which they attributed to CSA-mediated inhibition of pyruvate dehydrogenase and reduction of mitochondrial pyruvate metabolism (Prabhu et al., 2014). Other studies have found that cysteine administration contributes to accumulation of intracellular reduced glutathione that triggers reductive stress (Kolossov et al., 2015; Singh et al., 2015; Zhang et al., 2012). Our data, on the other hand, show that cysteine susceptibility does not depend on any particular cysteine metabolic pathway but rather is a feature of cysteine’s ability to alter the intracellular redox state. Indeed, we did not find evidence of accumulation of cysteine byproducts in our cells after labeling with ^13^C3-L-cysteine. Additionally, we found immediate depletion of intracellular NADPH and reduced glutathione, arguing that intracellular cysteine in these cells consumes rather than produces glutathione. Our findings that mutation of two reactive cysteine residues in Trx2, cysteine 90 and 93, abrogate its rescue of NAC-induced H_2_O_2_ production suggest that reactive cysteine residues on redox enzymes are required for cysteine-induced reductive stress. Therefore, it is likely that intracellular cysteine accumulation drives a reductive phenotype that overwhelms electron acceptors. This phenomenon is similar to the susceptibility of cysteine residues on numerous metabolic enzymes to the redox state of the cell. For example, several enzymes involved in glucose and amino acid metabolism, including pyruvate dehydrogenase, mitochondrial pyruvate carrier, aconitase, and phosphoglycerate dehydrogenase, have all been shown to be inhibited by oxidation of reactive cysteine residues (Halestrap, 1976; Han et al., 2005; Hurd et al., 2012; Mullarky et al., 2016). In our studies, metabolic tracing experiments suggest that the effects of NAC on these pathways is minimal and does not adequately explain the rapid and severe reduction in mitochondrial metabolism. Therefore, it is likely that the predominant mechanism of cysteine toxicity in our cells is through mitochondrial reductive stress that drives H_2_O_2_ production.

Our results also demonstrate that cysteine susceptibility synergizes with glucose starvation in GBM cells, providing therapeutic avenues to leverage synergistic regimens. Because detoxification of mitochondrial H_2_O_2_ depends on NADPH availability, glucose starvation depletes pentose phosphate-derived NADPH, thereby potentiating cysteine-induced toxicity. NAC is FDA-approved for several indications, including acetaminophen toxicity, contrast-induced nephropathy, and as an inhalational asthma treatment (Conrad et al., 2015; Smilkstein et al., 1988; Xu et al., 2016). It has been tested as a treatment, either alone or in combination with other cysteine-promoting drugs, to prevent oxidative stress that occurs after hypoxic-ischemic brain damage in stroke (Karuppagounder et al., 2018; Krzyzanowska et al., 2016; Krzyzanowska et al., 2017; Sabetghadam et al., 2020). Glucose starvation strategies, from anti-hyperglyemic drugs to the ketogenic diet, have also been tested alone and in combination with putative synergistic regimens for cancer treatment (Hopkins et al., 2018). Therefore, rational combination of cysteine compounds with glucose-lowering treatments may enhance cysteine toxicity in GBM and spare normal cells since primary human fetal astrocytes were unaffected by this regimen.

Several important questions remain unanswered from this work. Why does GBM express higher levels of cysteine and methionine metabolic genes than every other cancer? One explanation may be that GBM depends to a greater extent on cystine uptake to release glutamate through Xc activity, causing excitotoxicity and death of surrounding cells that supports expansion within the skull case (Noch and Khalili, 2009). GBM cells might also be dependent on cystine uptake to fuel glutathione production, which is especially important to mitigate the effects of ROS production under hypoxia (Chung et al., 2005; Ogunrinu and Sontheimer, 2010). The gasotransmitter hydrogen sulfide (H_2_S) is formed from cysteine-derived 3-mercaptopyruvate and serves as a potent activator of mitochondrial ATP production at low levels and inhibitor at high levels (Bonifacio et al., 2021). Though the role of cysteine-derived H_2_S in mediating GBM growth and metabolic fitness remains controversial (Bronowicka-Adamska et al., 2017; Silver et al., 2021; Wrobel et al., 2014; Zhao et al., 2015), GBM cells may depend on sufficient cystine uptake to maintain H_2_S-mediated energy production.

It is also unclear why GBM expresses lower levels of several mitochondrial redox enzymes than normal brain and several other cancers. Prior studies have shown that the brain may produce less ROS than other organs in response to ROS-inducing compounds and during aging (Tokumaru et al., 1996). Therefore, GBM may not require comparably high levels of redox enzyme expression as other organs. It is also possible that high levels of intracellular cysteine compensate for reduced redox enzyme expression and are sufficient to detoxify ROS associated with GBM growth. Because therapeutic resistance to the alkylating agent temozolomide involves genomic instability and hypermutation (Choi et al., 2018; Salazar-Ramiro et al., 2016; Yu et al., 2021), GBM may also require low levels of redox enzyme activity to maintain a high ROS burden. Fortunately, this simultaneous cystine avidity and reduced redox enzyme expression provide a unique opportunity to leverage susceptibility to reductive stress using cysteine-promoting compounds for GBM treatment.

In summary, we have shown that GBM possesses a unique susceptibility to the amino acid cysteine, which induces mitochondrial reductive stress that drives toxic accumulation of mitochondrial H_2_O_2_. This susceptibility is exacerbated by glucose starvation, making strategies to lower glucose levels synergistic with cysteine-promoting therapy. Future studies will validate these compounds to leverage cysteine susceptibility in animal models of GBM.

## STAR Methods

### Cell culture

667 and 603 glioma cells were obtained from Cameron Brennan at Memorial Sloan Kettering Cancer Center. 667 cells were cultured in 1:1 Neurobasal medium and DMEM/F12 medium supplemented with Glutamax, HEPES, sodium pyruvate, minimal essential amino acids, Pen/Strep, B27 supplement minus vitamin A, heparin (2 μg/ml), and EGF and FGF (20 μg/ml) (all from Life Technologies, Waltham, MA, USA). 603 cells were cultured in the same medium as 667 cells except with the addition of 20 ng/ml PDGF-AA and PDGF-BB (Shenandoah). Primary human fetal astrocytes were purchased from Thermo Fisher (NC9711462) and cultured in DMEM with 15% FBS. A549 (ATCC Cat# A549, RRID:CVCL_0023), MCF7 (ATCC Cat# MCF7, RRID:CVCL_0031), H1975 (ATCC Cat# CRL-5908, RRID:CVCL_1511), HT-29 (ATCC Cat# HT-29, RRID:CVCL_0320), and HPAF-II (ATCC Cat# CRL-1997, RRID:CVCL_0313) cells were purchased from ATCC. A549 and HT-29 cells were cultured in DMEM with 10% FBS and Pen-strep, MCF7 and HPAF-II cells were cultured in MEM with 10% FBS and Pen-strep, and H1975 cells were cultured in RPMI medium with 10% FBS and Pen-strep. Cell cultures were maintained at 37□°C under 5% CO2.

### Chemicals and Reagents

The following chemicals were purchased from Sigma: N-acetylcysteine (A7250), L-cysteine (168149), L-methionine (M5308), 2-deoxyglucose (2-DG) (D6134), D-cysteine (30095), L-cystine (C6727), glutathione reduced ethyl ester (GREE) (G1404), reduced glutathione (GSH) (G6013), oxidized glutathione (GSSG) (G4376), catalase (C1345), pCMB (C5913), and H_2_O_2_ (H1009). U-13C6-D-glucose (CLM-1396) and 13C3-L-cysteine (CLM-4320-H) were purchased from Cambridge Isotope Laboratories. Recombinant human Trx1 (LFP0001) and TrxR2 (LFP0019) were purchased from Life Technologies.

### Vectors

Human *TXN*, *TXN2*, *GSR*, *TXNRD2*, and *CAT* expression vectors were purchased from Genscript and cloned into the pLenti-Puro vector within BamHI and EcoRI restriction sites. The 3X-Myc-EGFP-OMP25 (Addgene plasmid number 83355) and 3X-HA-EGFP-OMP25 (Addgene plasmid number 83356) vectors were purchased from Addgene and cloned into the pLenti-Puro vector within BamHI and EcoRI restriction sites. The Trx2 C90S/C93S mutant was prepared from *TXN2* using overlap PCR with an oligonucleotide containing the desired mutation.

### Western blot

Whole-cell lysates were prepared in CST buffer (Cell Signaling Technology) containing protease inhibitor (Sigma) and were rotated at 4°C for 30 minutes before DNA was pelleted. Samples (60 μg) were mixed with 50 mM DTT (Thermo Scientific) and loaded on 4-12% Tris-Glycine gels (Thermo-Fisher) for SDS-PAGE before being transferred onto 0.2 μm or 0.45 μm nitrocellulose membrane with wet transfer cells (Bio-Rad Laboratories). After 30 minutes of blocking with 10% milk, blots were incubated with primary antibodies diluted to 1:5000 either at room temperature for 2 hours or overnight at 4°C. Blots were then incubated in either mouse or rabbit secondary antibody for 1 hour at room temperature before being developed using Dura or Femto enhanced chemiluminescence (Pierce) and imaged with ChemiDoc XRS+ and ImageLab Software version 6.1 (both from Bio-Rad Laboratories). The following primary antibodies were used: Cytochrome C (Abcam, ab13575, RRID:AB_300470), Trx1 (Proteintech, 14999-1-AP, RRID:AB_2272597), Trx2 (Cell Signaling Technology, 14907S, RRID:AB_2798645), TrxR1 (Cell Signaling Technology, 15140S, RRID:AB_2798725), TrxR2 (Cell Signaling Technology, 12029, RRID:AB_2797803), GR (Abcam, ab137513, RRID:AB_2732913), Citrate synthase (Cell Signaling Technology, 14309S, RRID:AB_2665545), Lamin A/C (Cell Signaling Technology, 4777S, RRID:AB_10545756) Cathepsin C (Santa Cruz Biotechnology, sc-74590, RRID:AB_2086955), p70 S6K (Cell Signaling Technology, 2708T, RRID:AB_390722), GM130 (BD Biosciences, 610823, RRID:AB_398142), Calnexin (Abcam, ab31290, RRID:AB_868628), beta-actin (Abcam, ab6276, RRID:AB_2223210).

### Cell proliferation assay

Cell proliferation was measured using the Cell Titer Glo reagent (Promega) according to the manufacturer’s instructions. Briefly, 10,000 cells were plated in a white-bottom 96-well plate. Drugs relevant to each experiment were added to corresponding wells, and the plate was placed in the incubator for 24-72 hours based on the specific experiment performed. Cells were incubated with Cell Titer Glo reagent for 10 minutes on a rocking platform, and luminescence was measured on a Synergy Neo 2 plate reader (BioTek Instruments).

### Oxygen consumption rate (OCR) measurement

Oxygen consumption rate was measured using the Seahorse XFe96 analyzer (Agilent) according to the manufacturer’s instructions. Briefly, 50,000 cells were plated in each well of a 96-well Seahorse plate. Adherent cells were plated the night prior to the assay, and non-adherent cells (603 and 667) were plated on Cell-Tak-coated (Corning) plates on the day of the assay according to the manufacturer’s instructions. At least 6 hours prior to the assay, the assay cartridge was hydrated in water in a non-CO_2_ incubator at 37°C. Cells were incubated in a non-CO_2_ incubator for 45 minutes. 10 mM NAC pH 7.4 was loaded into appropriate wells of the flux pack and placed into the XFe96 analyzer. The plate containing cells was subsequently loaded into the XFe96 analyzer, and OCR was analyzed using Seahorse Wave software.

### Mitochondrial Isolation

Mitochondrial isolation was performed as described previously (Chen et al., 2017). Briefly, cells were transduced with lentivirus expressing either 3X-Myc-EGFP-OMP25 or 3X-HA-EGFP-OMP25 and selected in Puromycin-containing medium for at least 1 week. 20-25 million cells were harvested, washed with KPBS, and subjected to homogenization in a glass homogenizer with 10 strokes without rotation. After homogenization, lysates were centrifuged at 2000×g for 2 min, and the supernatant was added to HA magnetic beads (Pierce) for mitochondrial immunoprecipitation followed by 3 washes in KPBS. Lysates were prepared by adding Triton X-100 buffer to the beads according to the protocol, and whole mitochondria were isolated by adding HA peptide (Thermo Fisher) to the beads and incubating the bead mixtures at 37°C for 10 minutes.

### Hydrogen peroxide measurement

Mitochondrial hydrogen peroxide levels were measured using the Amplex Red reagent (Thermo Fisher) according to the manufacturer’s instructions. Briefly, whole mitochondria were isolated using the method described above. 400,000 whole-cell equivalents were added to each well of a 96-well plate. Mitochondria were permeabilized with 2 μg/mL Alamethicin (Sigma, A4665), and either vehicle or 100 mM NAC pH 7.4 was added to the wells. The Amplex Red reagent, mixed with 200 mU/mL horseradish peroxidase (Thermo Fisher, 31491), was added to each well, and fluorescence was read at an excitation wavelength of 544 nm and emission wavelength of 590 nm for 45 minutes at room temperature on a Synergy Neo 2 plate reader (BioTek Instruments).

### Mitochondrial membrane potential measurement

Mitochondrial membrane potential was measured using the TMRE kit (Abcam) according to the manufacturer’s instructions. Briefly, 200,000 cells were treated with 10 mM NAC for the indicated time periods followed by incubation in 500 nM TMRE reagent. Fluorescence was read at an excitation wavelength of 544 nm and emission wavelength of 590 nm for 30 minutes at room temperature on a Synergy Neo 2 plate reader (BioTek Instruments).

### Glutathione measurement

Glutathione was measured using the Glutathione Glo reagent (Promega) according to the manufacturer’s instructions. Briefly, 20,000 cells were plated in a 96-well plate, treated with 10 mM NAC pH 7.4 for the indicated time periods, and incubated with either Total Glutathione Lysis Reagent or Oxidized Glutathione Lysis Reagent before luminescence was read on a Synergy Neo 2 plate reader (BioTek Instruments).

### NADPH measurement

NADPH was measured using the NADPH assay kit (Abcam) according to the manufacturer’s instructions. Briefly, 2 million cells were treated with either vehicle or 10 mM NAC. 6 replicates were used for each condition. At 5 minutes after treatment, cells were extracted in NADP/NADPH extraction buffer, and samples were deproteinated using 10 kD spin columns (ab93349). Total NADP/NADPH and NADPH levels were measured as per the protocol, and readings were taken 1 hour after NADPH reaction mix incubation at OD450 nm on a Synergy Neo 2 plate reader (BioTek Instruments).

### Apoptosis and necrosis measurement

Apoptosis and necrosis were measured using the apoptosis/necrosis detection kit (Enzo). Briefly, 200,000 cells were treated with 10 mM NAC pH 7.4 for the indicated time periods before being incubated with the apoptosis (Annexin V) and necrosis (7-AAD) detection reagents according to the protocol. Fluorescence was detected on an Attune (BD Biosciences) with software.

### Caspase measurement

Caspase activity measurements were made using the Caspase assay kit (Abcam) according to the manufacturer’s instructions. Briefly, 2 million cells were treated with either vehicle or 10 mM NAC pH 7.4. Caspase 3 assay solution was added to each well, and fluorescence was measured at excitation wavelength of 544 nm and emission wavelength of 620 nm on a Synergy Neo 2 plate reader (BioTek Instruments).

### Mitochondrial EM

One million 667 cells in suspension were washed once with PBS and then fixed *in vitro* with a modified Karnovsky’s fix (Ito and Karnovsky, 1968) and a secondary fixation in reduced osmium tetroxide (De Bruijn, 1973). Following dehydration, the monolayers were embedded in an epon analog resin. *En face* ultrathin sections (65 nm) were contrasted with lead citrate (Venable and Coggeshall, 1965) and viewed on a JEM 1400 electron microscope (JEOL, USA, Inc., Peabody, MA) operated at 100 kV. Digital images were captured on a Veleta 2K × 2K CCD camera (EMSIS, Muenster, Germany). The number of visible cristae per mitochondria and total mitochondria were quantified from 312.5 μm^2^ area.

### Cytochrome C release assay

Cytosolic Cytochrome C release from the mitochondria was measured by immunoblot of cytoplasmic and mitochondrial lysates. Briefly, whole mitochondria were isolated as above and lysed in 1X CST buffer from HA magnetic beads. Cytoplasmic proteins were isolated by collecting supernatant after incubation of cellular homogenates with HA magnetic beads. The cytoplasmic fraction was cleared with 25 μl additional HA magnetic beads for 3.5 minutes with rotation at 4°C to further purify the cytoplasmic fraction. Lysates were prepared by adding 10X CST containing protease inhibitor to this fraction. Mitochondrial and cytoplasmic fractions were subjected to SDS-PAGE followed by immunoblot with Cytochrome C antibody (Abcam) at 1:500 dilution.

### Confocal Microscopy

For confocal microscopy of 667 cells, cells were plated on PDL-coated coverslips and centrifuged at 200×g for 2 minutes. Cells were incubated with 100 nM Mitotracker and 1 μg/mL Hoechst stain, and imaged using a Zeiss LSM990 microscope fit with a camera and analyzed with Zeiss software.

For mitochondrial hydrogen peroxide microscopy, 200,000 667 cells were plated on PDL-coated glass coverslips in PBS and attached by centrifugation at 200×g for 1 minute. 100 nM Mitotracker Red (Thermo), 1 μg/mL Hoechst stain, and 10 μM Mito-PY1 were added to each well and incubated at 37°C for 30 minutes. Cells were washed once in fresh PBS, and 10 mM NAC or vehicle was added to each well and incubated for 2 hours at 37°C.

### Metabolomics analysis

Metabolites were extracted using pre-cooled 80% methanol. Samples were then centrifuged at 4°C for 15 minutes at 14,000 rpm. The supernatants containing polar metabolites were dried down using a SpeedVac. Targeted LC/MS analyses were performed on a Q Exactive Orbitrap mass spectrometer (Thermo Scientific) coupled to a Vanquish UPLC system (Thermo Scientific). The Q Exactive operated in polarity-switching mode. A Sequant ZIC-HILIC column (2.1 mm i.d. × 150 mm, Merck) was used for separation of metabolites. Flow rate was set at 150 μL/min. Buffers consisted of 100% acetonitrile for mobile B, and 0.1% NH_4_OH/20 mM CH_3_COONH_4_ in water for mobile A. Gradient ran from 85% to 30% B in 20 min followed by a wash with 30% B and re-equilibration at 85% B. Data analysis was done using El-MAVEN (v0.12.0). Metabolites were identified based on exact mass within 5 ppm and standard retention times.

### Cysteine and methionine gene expression analysis

RNAseq-based gene expression profiles normalized as Fragments Per Kilobase of transcript per Million mapped reads (FPKMs) were downloaded from the Genomic Data Commons (GDC) Data Portal (https://portal.gdc.cancer.gov). These FPKMs were quantified from the BAMs re-aligned to the GRCh38 genome build and re-processed using the GDC’s standardized pipelines (https://gdc.cancer.gov/about-data/gdc-data-harmonization). The cysteine methionine metabolic signature genes were obtained from the Molecular Signature Database (MSigDB) Gene Set KEGG_CYSTEINE_AND_METHIONINE_METABOLISM (Subramanian et al., 2005). Single Sample Gene Set Enrichment Analysis (ssGSEA) was applied to score the metabolic signature in each tumor sample using their RNAseq expression profiles (Barbie et al., 2009). ssGSEA was implemented using the ‘gsva’ package (v1.30.0) in R.

### Differential proteomics analysis

The samples in this study comprise data from the National Cancer Institute’s Clinical Proteomic Tumor Analysis Consortium (CPTAC) (Edwards et al., 2015; Ellis et al., 2013). We downloaded the LC-MS/MS protein-quantitation data and associated clinical data of CPTAC’s 6 Discovery Studies, each of which characterized one of the following cancer types: clear cell renal cell carcinoma (ccRCC) (Clark et al., 2019), GBM (Wang et al., 2021), lung adenocarcinoma (LUAD) (Gillette et al., 2020), lung squamous cell carcinoma (LSCC) (Satpathy et al., 2021), and uterine corpus endometrial carcinoma (UCEC) (Dou et al., 2020). We used base R functions (R-Core-Team, 2020) and dplyr (Wickham et al., 2021) to isolate the spectral counts for only those peptides which uniquely mapped to individual proteins. We then generated two unsupervised hierarchically clustered heatmaps for each cancer with clinical data annotations, one for redox enzyme expression and another for the expression of cysteine- and methionine-metabolizing proteins, using the R package ComplexHeatmap (Gu et al., 2016). We generated a list of redox proteins based on known redox enzymes compiled by surveying publicly available resources (https://www.rndsystems.com/research-area/redox-enzymes) and added glutathione reductase given its role in reductive stress.

Within each cancer type’s dataset, we also used base R functions and dplyr to calculate the ratio of each redox enzyme’s median expression in cancer tissues to its median expression in normal samples. These ratios were visualized using the ggplot2 and ggforce (Pedersen, 2021).

### Differential metabolomics analysis

Raw metabolomics ion counts were taken from Chinnaiyan et al. (Chinnaiyan et al., 2012) The original dataset included 69 samples from patients with glioma WHO grade II (n=18), III (n=18), and IV (n=33). Metabolite abundances were first corrected for run day batch effects using median scaling, normalized using the probabilistic quotient normalization (Dieterle et al., 2006) using metabolites with less than 20% missing values to generate the reference sample, and log_2_ transformed. Twelve metabolites annotated with the pathway “Cysteine, methionine, SAM, taurine metabolism” (2-hydroxybutyrate, cystathionine, cysteine, cysteine sulfinic acid, cystine, homocysteine, hypotaurine, methionine, N-acetylmethionine, S-adenosylhomocysteine, S-methylcysteine, taurine) were considered for statistical analysis. Associations between metabolite abundance and tumor grade were estimated using Kendall’s correlation, and p-values were corrected for multiple testing using the Benjamini-Hochberg method for controlling the false discovery rate (Hochberg, 1995). Metabolites with adjusted p-values less than 0.05 were considered significant. The repository of the R code to reproduce the metabolomics results can be found at https://github.com/krumsieklab/glioma-cysteine.

### Statistical Analysis

Statistical analyses were conducted using GraphPad Prism version 8.0 (La Jolla, CA, USA). Data are mean +/− standard deviation unless otherwise stated. t-tests were used to measure significance between 2 samples, and ANOVA was used to measure significance between 3 or more groups, with results being adjusted for multiple comparisons. All error bars indicate the mean +/− standard deviation unless otherwise specified.

## Data and Software Availability

All original data from this manuscript are available here:

## Inclusion and Diversity

One or more of the authors of this paper self-identifies as an underrepresented ethnic minority in science. One or more of the authors of this paper self-identifies as a member of the LGBTQ+ community. One or more of the authors of this paper received support from a program designed to increase minority representation in science.

## Materials Availability

All unique/stable reagents generated in this study are available from the Lead Contact without restriction.

## Lead Contact Statement

Further information and requests for resources and reagents should be directed to and will be fulfilled by the Lead Contact, Evan Noch.

---

**Figure S1.**
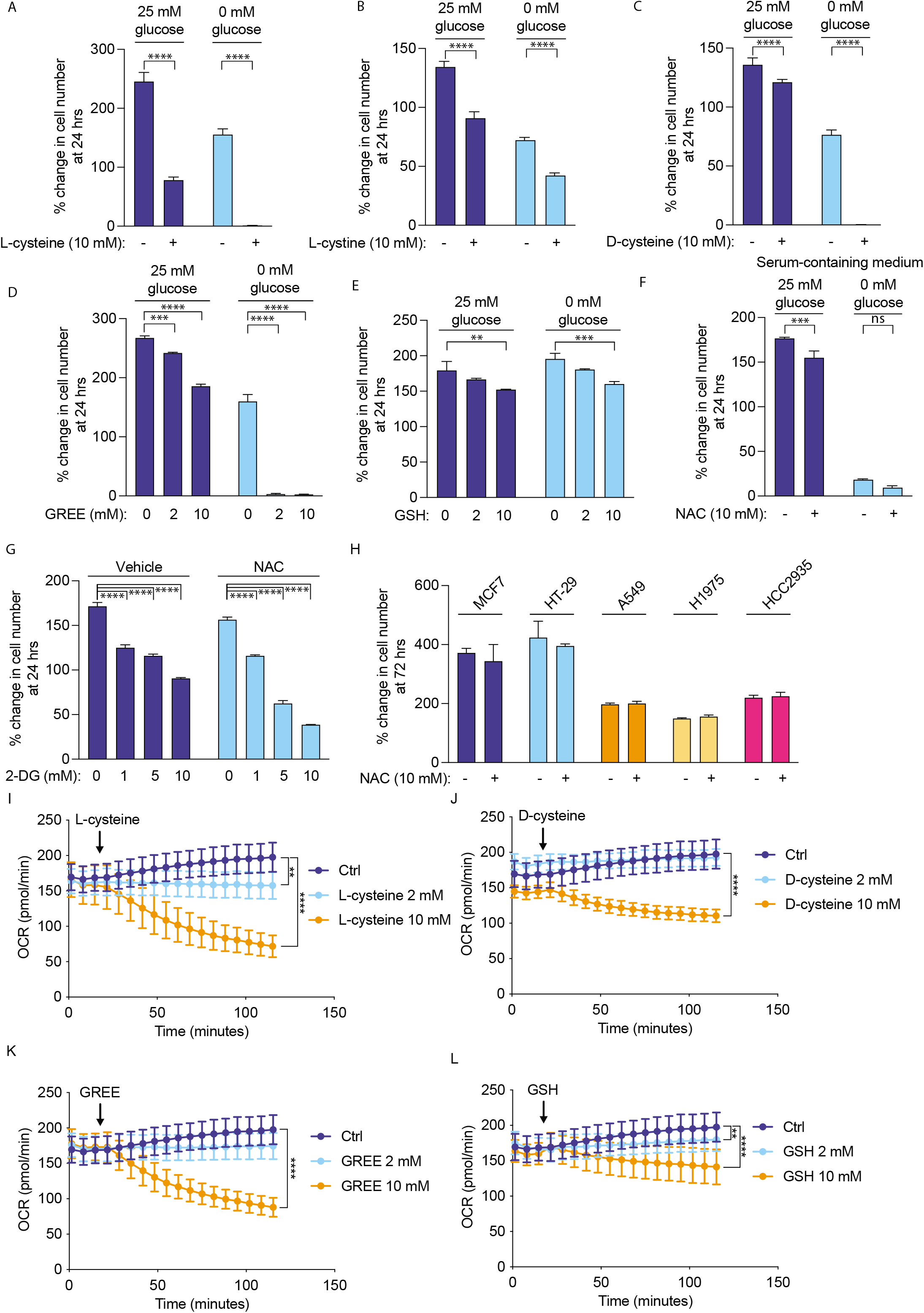
Cysteine-containing compounds suppress growth and reduce oxygen consumption in GBM cells. 667 cells were treated with 10 mM L-cysteine (A), 10 mM L-cystine (B), 10 mM D-cysteine (C), 2 or 10 mM glutathione reduced ethyl ester (GREE) (D), or reduced glutathione (GSH) (E) under normal glucose conditions (25 mM) or glucose starvation (0 mM). Growth was measured at 24 hours. (F) 667 cells were grown in DMEM containing 10% FBS and were treated with 10 mM NAC under normal glucose conditions (25 mM) or glucose starvation, and growth was measured at 24 hours. (G) 667 cells were treated with 1, 5, or 10 mM 2-deoxyglucose (2-DG) along with vehicle or 10 mM NAC, and growth was measured at 24 hours. (H) MCF7, HT-29, A549, H1975, and HCC2935 cells were treated with 10 mM NAC under normal glucose conditions (25 mM), and growth was measured at 72 hours. 667 cells were treated with 2 or 10 mM L-cysteine (I) 2 or 10 mM D-cysteine (J), 2 or 10 mM GREE (K), or 2 or 10 mM GSH (L), and oxygen consumption rate was measured using the Seahorse assay. *, p < 0.05, **, p < 0.01, ***, p < 0.001, ****, p < 0.0001.

**Figure S2.**
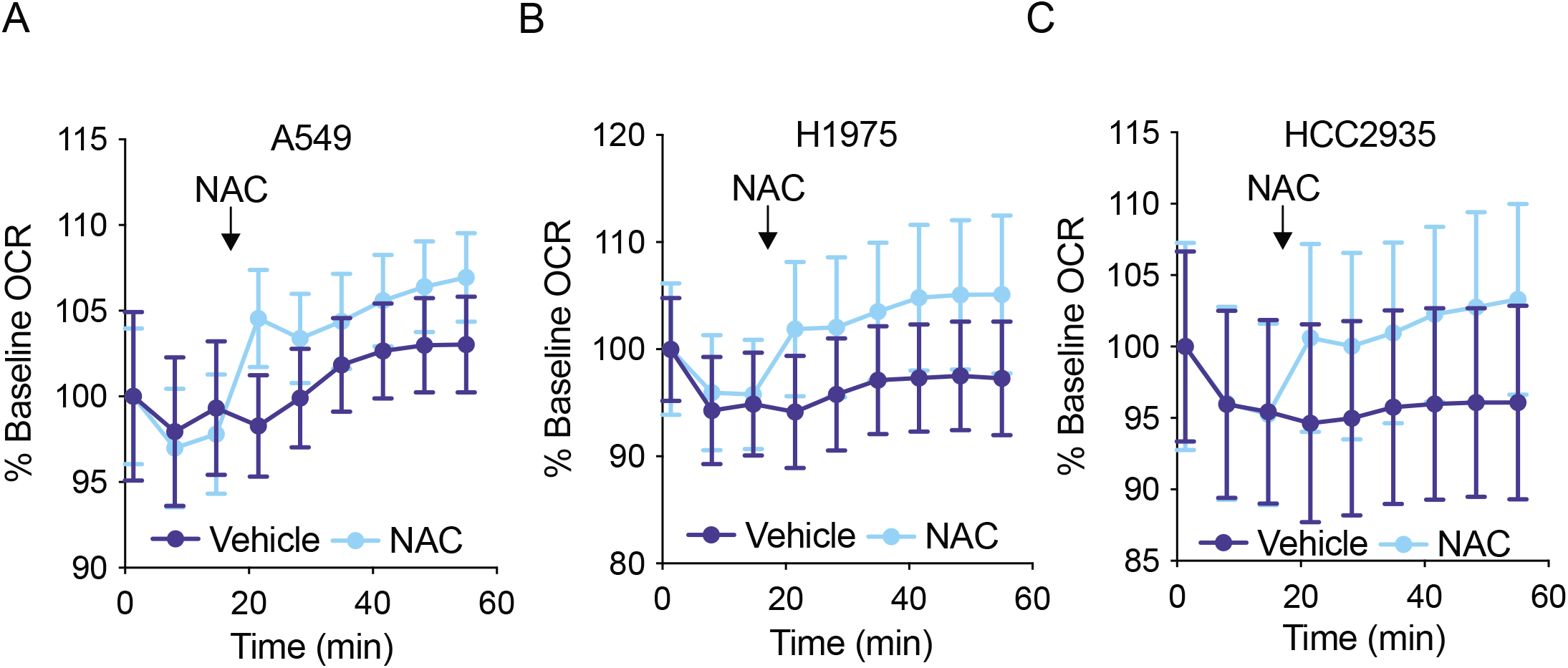
NAC does not reduce oxygen consumption in non-glioma cells. A549 (A), MDAMB231 (B), MDAMB468 (C), H1975 (D), and HCC2935 (E) cells were treated with 10 mM NAC, and oxygen consumption was measured with the Seahorse assay.

**Figure S3.**
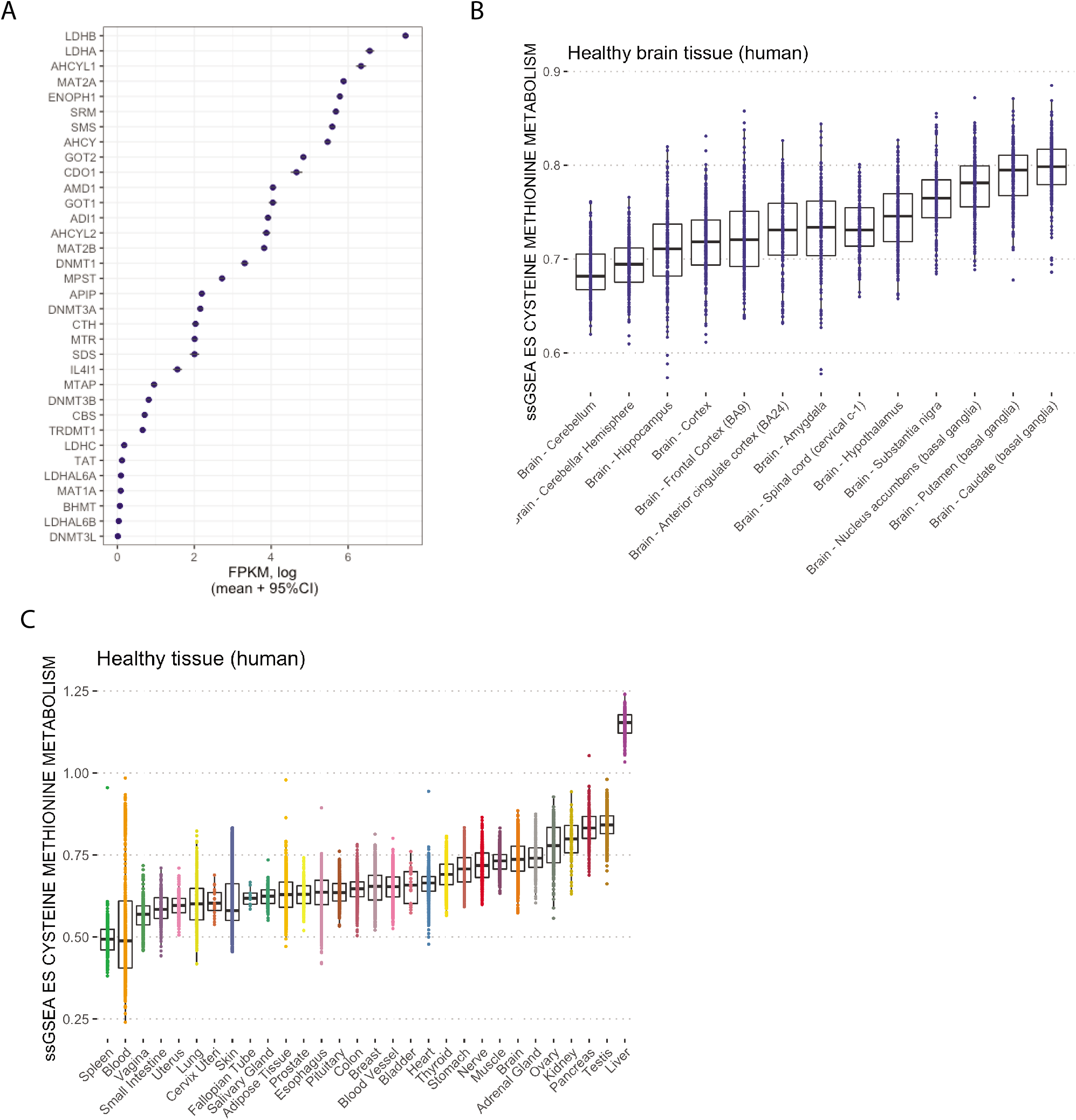
Distribution of cysteine and methionine gene signature in healthy tissues and normal brain. Distribution of cysteine and methionine KEGG genes across TCGA GBM samples (A), normal brain regions (B), and normal human tissues (C).

**Figure S4.**
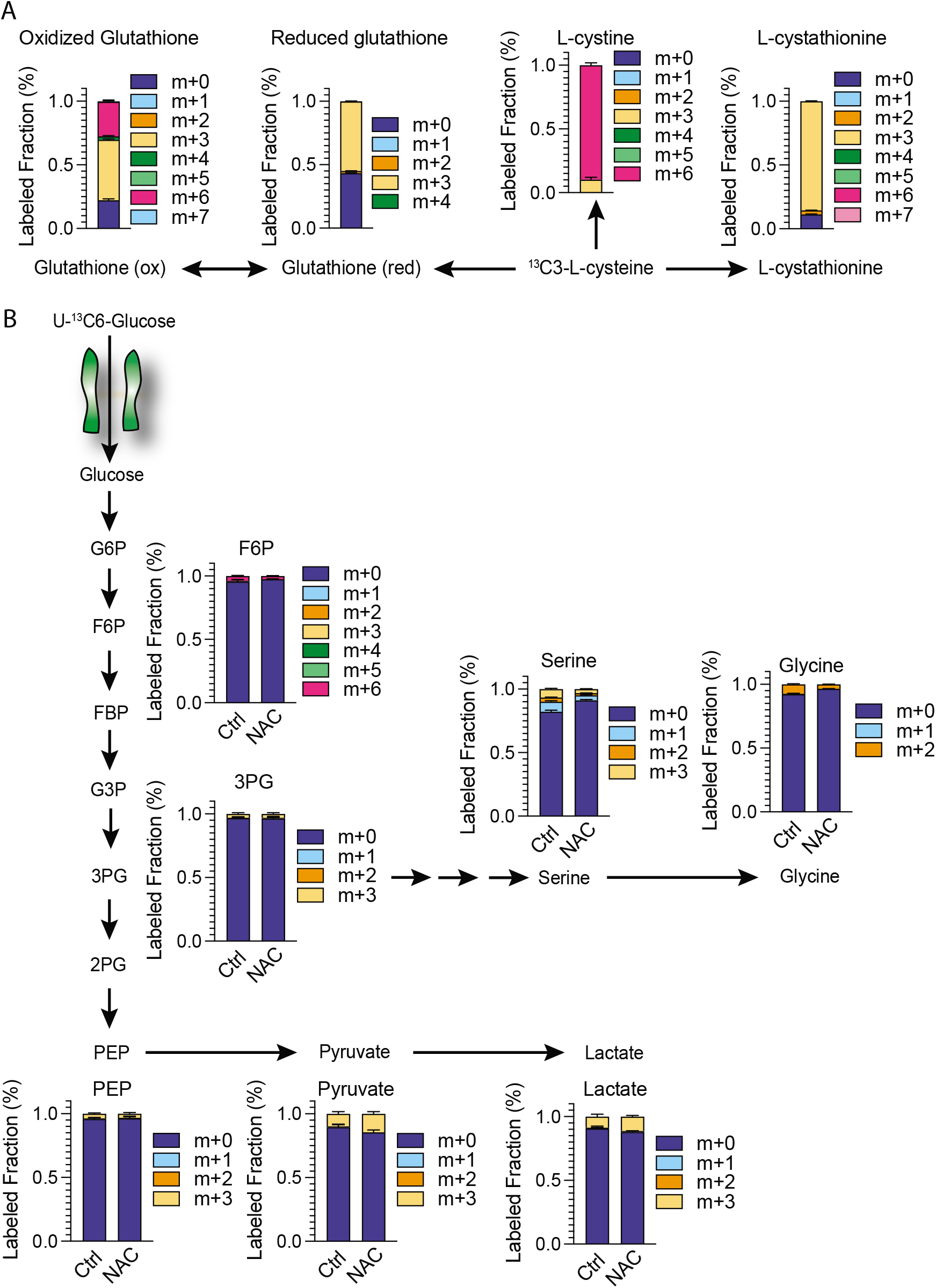
^13^C3-L-Cysteine and U-^13^C6-glucose metabolic flux in GBM cells. (A) 667 cells were treated with 10 mM ^13^C3-L-cysteine for 5 minutes, and labeled metabolites were measured by LC/MS. (B) GBM cells were treated with 10 mM ^13^C6-glucose and 10 mM NAC for 5 minutes, and labeled metabolites were measured by LC/MS.

**Figure S5.**
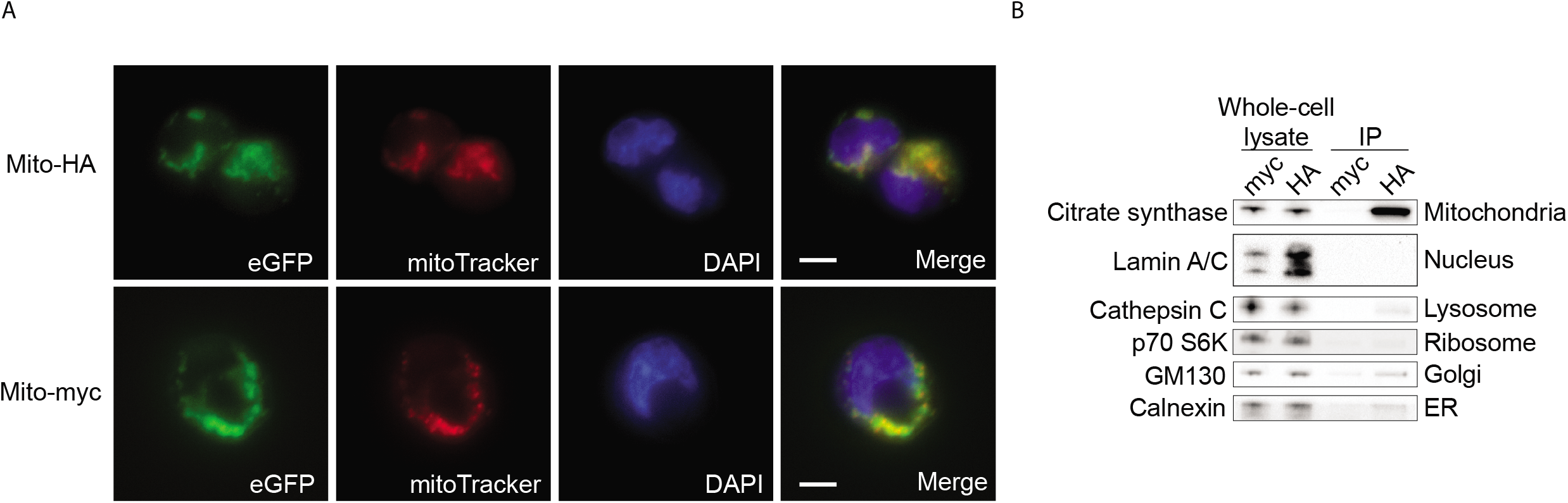
Immunoprecipitation of 667 mitochondria using an HA-tag based method. (A) 667 cells expressing HA-OMP25-EGFP or Myc-OMP25-EGFP were incubated with 100 nM Mitotracker Deep Red and 1 μg/mL Hoechst stain, followed by confocal microscopy. Scale bars are 10 μm. (B) 667 cells expressing HA-OMP25-EGFP or Myc-OMP25-EGFP were immunoprecipitated using HA magnetic beads, and lysates were run on SDS-PAGE along with whole-cell lysates prepared before immunoprecipitation. The indicated cell compartment-specific antibodies were used.

**Figure S6.**
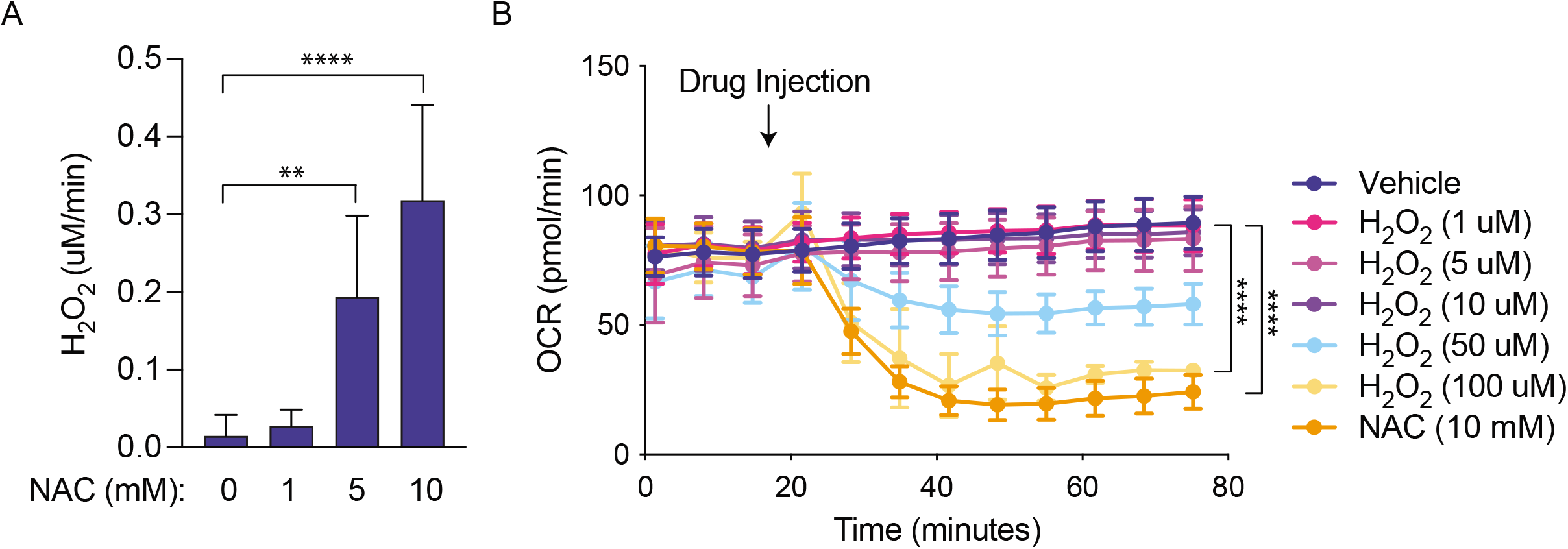
GBM cells induce greater OCR reduction than 100 μM H_2_O_2_. (A) 1, 5, or 10 mM NAC was added to mitochondrial assay buffer, and H_2_O_2_ production was measured with Amplex Red reagent. (B) 10 mM NAC or 1, 5, 10, 50, or 100 μM H_2_O_2_ was added to 667 cells, and oxygen consumption was measured with the Seahorse assay. **, p < 0.01, ****, p < 0.0001.

**Figure S7.**
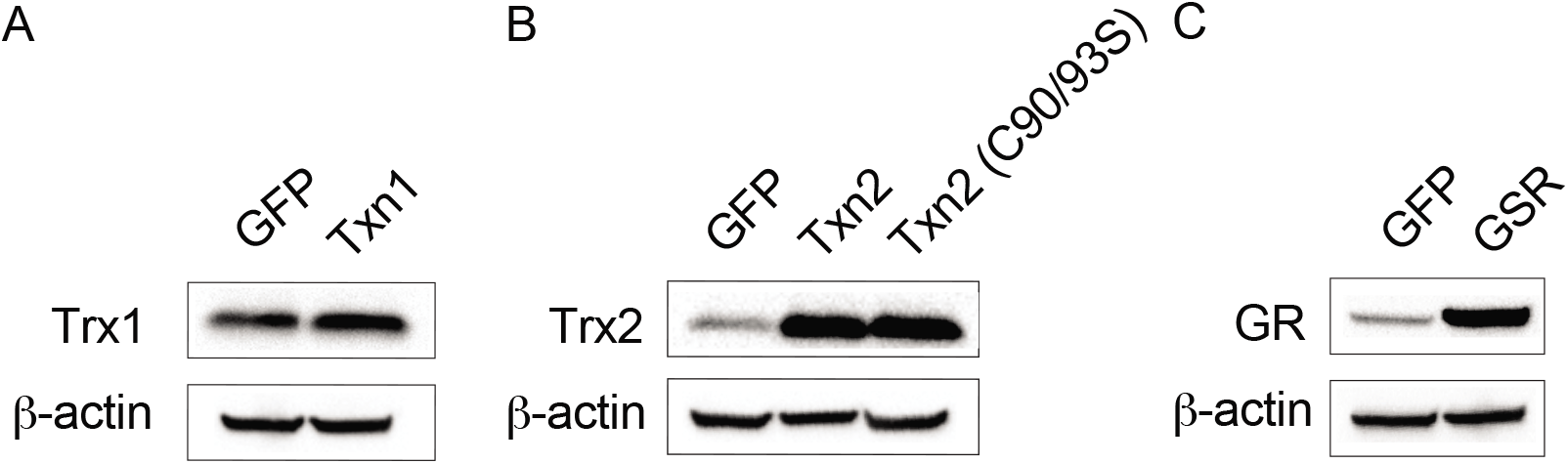
Genetic overexpression of redox enzymes in 667 cells. 667 cells were transduced with lentivirus expressing GFP or Txn1 (A), Txn2 or Txn2 (C90/93S) (B), or GSR (C). Whole-cell lysates were subjected to immunoblot with the indicated antibodies.

## References

Banjac, A., Perisic, T., Sato, H., Seiler, A., Bannai, S., Weiss, N., Kolle, P., Tschoep, K., Issels, R.D., Daniel, P.T., et al. (2008). The cystine/cysteine cycle: a redox cycle regulating susceptibility versus resistance to cell death. Oncogene 27, 1618–1628.

Barbie, D.A., Tamayo, P., Boehm, J.S., Kim, S.Y., Moody, S.E., Dunn, I.F., Schinzel, A.C., Sandy, P., Meylan, E., Scholl, C., et al. (2009). Systematic RNA interference reveals that oncogenic KRAS-driven cancers require TBK1. Nature 462, 108–112.

Barthel, A., Okino, S.T., Liao, J., Nakatani, K., Li, J., Whitlock, J.P., Jr., and Roth, R.A. (1999). Regulation of GLUT1 gene transcription by the serine/threonine kinase Akt1. J Biol Chem 274, 20281–20286.

Bavarsad Shahripour, R., Harrigan, M.R., and Alexandrov, A.V. (2014). N-acetylcysteine (NAC) in neurological disorders: mechanisms of action and therapeutic opportunities. Brain Behav 4, 108–122.

Bonifacio, V.D.B., Pereira, S.A., Serpa, J., and Vicente, J.B. (2021). Cysteine metabolic circuitries: druggable targets in cancer. Br J Cancer 124, 862–879.

Briggs, K.J., Koivunen, P., Cao, S., Backus, K.M., Olenchock, B.A., Patel, H., Zhang, Q., Signoretti, S., Gerfen, G.J., Richardson, A.L., et al. (2016). Paracrine Induction of HIF by Glutamate in Breast Cancer: EglN1 Senses Cysteine. Cell 166, 126–139.

Bronowicka-Adamska, P., Bentke, A., and Wrobel, M. (2017). Hydrogen sulfide generation from l-cysteine in the human glioblastoma-astrocytoma U-87 MG and neuroblastoma SHSY5Y cell lines. Acta Biochim Pol 64, 171–176.

Chen, W.W., Freinkman, E., and Sabatini, D.M. (2017). Rapid immunopurification of mitochondria for metabolite profiling and absolute quantification of matrix metabolites. Nat Protoc 12, 2215–2231.

Chinnaiyan, P., Kensicki, E., Bloom, G., Prabhu, A., Sarcar, B., Kahali, S., Eschrich, S., Qu, X., Forsyth, P., and Gillies, R. (2012). The metabolomic signature of malignant glioma reflects accelerated anabolic metabolism. Cancer Res 72, 5878–5888.

Choi, S., Yu, Y., Grimmer, M.R., Wahl, M., Chang, S.M., and Costello, J.F. (2018). Temozolomide-associated hypermutation in gliomas. Neuro Oncol 20, 1300–1309.

Chung, W.J., Lyons, S.A., Nelson, G.M., Hamza, H., Gladson, C.L., Gillespie, G.Y., and Sontheimer, H. (2005). Inhibition of cystine uptake disrupts the growth of primary brain tumors. J Neurosci 25, 7101–7110.

Clark, D.J., Dhanasekaran, S.M., Petralia, F., Pan, J., Song, X., Hu, Y., da Veiga Leprevost, F., Reva, B., Lih, T.-S.M., Chang, H.-Y., et al. (2019). Integrated Proteogenomic Characterization of Clear Cell Renal Cell Carcinoma. Cell 179, 964–983.e931.

Combs, J.A., and DeNicola, G.M. (2019). The Non-Essential Amino Acid Cysteine Becomes Essential for Tumor Proliferation and Survival. Cancers (Basel) 11.

Conrad, C., Lymp, J., Thompson, V., Dunn, C., Davies, Z., Chatfield, B., Nichols, D., Clancy, J., Vender, R., Egan, M.E., et al. (2015). Long-term treatment with oral N-acetylcysteine: affects lung function but not sputum inflammation in cystic fibrosis subjects. A phase II randomized placebo-controlled trial. J Cyst Fibros 14, 219–227.

De Bruijn, W. (1973). Glycogen its chemistry and morphologic appearance in the electron microscope. J. Ultrastruct. Res. 42, 29–50.

Dieterle, F., Ross, A., Schlotterbeck, G., and Senn, H. (2006). Probabilistic quotient normalization as robust method to account for dilution of complex biological mixtures. Application in 1H NMR metabonomics. Anal Chem 78, 4281–4290.

Dou, Y., Kawaler, E.A., Cui Zhou, D., Gritsenko, M.A., Huang, C., Blumenberg, L., Karpova, A., Petyuk, V.A., Savage, S.R., Satpathy, S., et al. (2020). Proteogenomic Characterization of Endometrial Carcinoma. Cell 180, 729–748.e726.

Edwards, N.J., Oberti, M., Thangudu, R.R., Cai, S., McGarvey, P.B., Jacob, S., Madhavan, S., and Ketchum, K.A. (2015). The CPTAC Data Portal: A Resource for Cancer Proteomics Research. J Proteome Res 14, 2707–2713.

Ellis, M.J., Gillette, M., Carr, S.A., Paulovich, A.G., Smith, R.D., Rodland, K.K., Townsend, R.R., Kinsinger, C., Mesri, M., Rodriguez, H., et al. (2013). Connecting genomic alterations to cancer biology with proteomics: the NCI Clinical Proteomic Tumor Analysis Consortium. Cancer Discov 3, 1108–1112.

Finn, N.A., and Kemp, M.L. (2012). Pro-oxidant and antioxidant effects of N-acetylcysteine regulate doxorubicin-induced NF-kappa B activity in leukemic cells. Mol Biosyst 8, 650–662.

Fu, L., Liu, K., Sun, M., Tian, C., Sun, R., Morales Betanzos, C., Tallman, K.A., Porter, N.A., Yang, Y., Guo, D., et al. (2017). Systematic and Quantitative Assessment of Hydrogen Peroxide Reactivity With Cysteines Across Human Proteomes. Mol Cell Proteomics 16, 1815–1828.

Gillette, M.A., Satpathy, S., Cao, S., Dhanasekaran, S.M., Vasaikar, S.V., Krug, K., Petralia, F., Li, Y., Liang, W.-W., Reva, B., et al. (2020). Proteogenomic Characterization Reveals Therapeutic Vulnerabilities in Lung Adenocarcinoma. Cell 182, 200–225.e235.

Gu, Z., Eils, R., and Schlesner, M. (2016). Complex heatmaps reveal patterns and correlations in multidimensional genomic data. Bioinformatics 32, 2847–2849.

Halestrap, A.P. (1976). The mechanism of the inhibition of the mitochondrial pyruvate transportater by alpha-cyanocinnamate derivatives. Biochem J 156, 181–183.

Han, D., Canali, R., Garcia, J., Aguilera, R., Gallaher, T.K., and Cadenas, E. (2005). Sites and mechanisms of aconitase inactivation by peroxynitrite: modulation by citrate and glutathione. Biochemistry 44, 11986–11996.

Hochberg, Y.B.a.Y. (1995). Controlling the False Discovery Rate: A Practical and Powerful Approach to Multiple Testing. Journal of the Royal Statistical Society 57, 289–300.

Hopkins, B.D., Pauli, C., Du, X., Wang, D.G., Li, X., Wu, D., Amadiume, S.C., Goncalves, M.D., Hodakoski, C., Lundquist, M.R., et al. (2018). Suppression of insulin feedback enhances the efficacy of PI3K inhibitors. Nature 560, 499–503.

Hurd, T.R., Collins, Y., Abakumova, I., Chouchani, E.T., Baranowski, B., Fearnley, I.M., Prime, T.A., Murphy, M.P., and James, A.M. (2012). Inactivation of pyruvate dehydrogenase kinase 2 by mitochondrial reactive oxygen species. J Biol Chem 287, 35153–35160.

Ilic, N., Birsoy, K., Aguirre, A.J., Kory, N., Pacold, M.E., Singh, S., Moody, S.E., DeAngelo, J.D., Spardy, N.A., Freinkman, E., et al. (2017). PIK3CA mutant tumors depend on oxoglutarate dehydrogenase. Proc Natl Acad Sci U S A 114, E3434–E3443.

Ito, S., and Karnovsky, M. (1968). Formaldehyde -glutaraldehyde fixatives containing trinitro compounds. J Cell Biol 39, 168A–169A.

Karuppagounder, S.S., Alin, L., Chen, Y., Brand, D., Bourassa, M.W., Dietrich, K., Wilkinson, C.M., Nadeau, C.A., Kumar, A., Perry, S., et al. (2018). N-acetylcysteine targets 5 lipoxygenase-derived, toxic lipids and can synergize with prostaglandin E2 to inhibit ferroptosis and improve outcomes following hemorrhagic stroke in mice. Ann Neurol 84, 854–872.

Kolossov, V.L., Beaudoin, J.N., Ponnuraj, N., DiLiberto, S.J., Hanafin, W.P., Kenis, P.J., and Gaskins, H.R. (2015). Thiol-based antioxidants elicit mitochondrial oxidation via respiratory complex III. Am J Physiol Cell Physiol 309, C81–91.

Koppula, P., Zhang, Y., Shi, J., Li, W., and Gan, B. (2017). The glutamate/cystine antiporter SLC7A11/xCT enhances cancer cell dependency on glucose by exporting glutamate. J Biol Chem 292, 14240–14249.

Korge, P., Calmettes, G., and Weiss, J.N. (2015). Increased reactive oxygen species production during reductive stress: The roles of mitochondrial glutathione and thioredoxin reductases. Biochim Biophys Acta 1847, 514–525.

Krzyzanowska, W., Pomierny, B., Budziszewska, B., Filip, M., and Pera, J. (2016). N-Acetylcysteine and Ceftriaxone as Preconditioning Strategies in Focal Brain Ischemia: Influence on Glutamate Transporters Expression. Neurotox Res 29, 539–550.

Krzyzanowska, W., Pomierny, B., Bystrowska, B., Pomierny-Chamiolo, L., Filip, M., Budziszewska, B., and Pera, J. (2017). Ceftriaxone- and N-acetylcysteine-induced brain tolerance to ischemia: Influence on glutamate levels in focal cerebral ischemia. PLoS One 12, e0186243.

Kuljanin, M., Mitchell, D.C., Schweppe, D.K., Gikandi, A.S., Nusinow, D.P., Bulloch, N.J., Vinogradova, E.V., Wilson, D.L., Kool, E.T., Mancias, J.D., et al. (2021). Reimagining high-throughput profiling of reactive cysteines for cell-based screening of large electrophile libraries. Nat Biotechnol 39, 630–641.

LeBoeuf, S.E., Wu, W.L., Karakousi, T.R., Karadal, B., Jackson, S.R., Davidson, S.M., Wong, K.K., Koralov, S.B., Sayin, V.I., and Papagiannakopoulos, T. (2020). Activation of Oxidative Stress Response in Cancer Generates a Druggable Dependency on Exogenous Non-essential Amino Acids. Cell Metab 31, 339–350 e334.

Lin, W., Wang, C., Liu, G., Bi, C., Wang, X., Zhou, Q., and Jin, H. (2020). SLC7A11/xCT in cancer: biological functions and therapeutic implications. Am J Cancer Res 10, 3106–3126.

Long, Y., Tao, H., Karachi, A., Grippin, A.J., Jin, L., Chang, Y.E., Zhang, W., Dyson, K.A., Hou, A.Y., Na, M., et al. (2019). Dysregulation of glutamate transport enhances Treg function that promotes VEGF blockade resistance in glioblastoma. Cancer Res.

Marin-Valencia, I., Yang, C., Mashimo, T., Cho, S., Baek, H., Yang, X.L., Rajagopalan, K.N., Maddie, M., Vemireddy, V., Zhao, Z., et al. (2012). Analysis of tumor metabolism reveals mitochondrial glucose oxidation in genetically diverse human glioblastomas in the mouse brain in vivo. Cell Metab 15, 827–837.

Mink, J.W., Blumenschine, R.J., and Adams, D.B. (1981). Ratio of central nervous system to body metabolism in vertebrates: its constancy and functional basis. Am J Physiol 241, R203–212.

Mullarky, E., Lucki, N.C., Beheshti Zavareh, R., Anglin, J.L., Gomes, A.P., Nicolay, B.N., Wong, J.C., Christen, S., Takahashi, H., Singh, P.K., et al. (2016). Identification of a small molecule inhibitor of 3-phosphoglycerate dehydrogenase to target serine biosynthesis in cancers. Proc Natl Acad Sci U S A 113, 1778–1783.

Noch, E., and Khalili, K. (2009). Molecular mechanisms of necrosis in glioblastoma: the role of glutamate excitotoxicity. Cancer Biol Ther 8, 1791–1797.

Ogunrinu, T.A., and Sontheimer, H. (2010). Hypoxia increases the dependence of glioma cells on glutathione. J Biol Chem 285, 37716–37724.

Pedersen, T. (2021). ggforce: Accelerating ‘ggplot2’. 0.3.3 ed2021.

Prabhu, A., Sarcar, B., Kahali, S., Yuan, Z., Johnson, J.J., Adam, K.P., Kensicki, E., and Chinnaiyan, P. (2014). Cysteine catabolism: a novel metabolic pathway contributing to glioblastoma growth. Cancer Res 74, 787–796.

R-Core-Team, R.C.T. (2020). R: A Language and Environment for Statistical Computing. R Foundation for Statistical Computing.

Robert, S.M., Buckingham, S.C., Campbell, S.L., Robel, S., Holt, K.T., Ogunrinu-Babarinde, T., Warren, P.P., White, D.M., Reid, M.A., Eschbacher, J.M., et al. (2015). SLC7A11 expression is associated with seizures and predicts poor survival in patients with malignant glioma. Sci Transl Med 7, 289ra286.

Sabetghadam, M., Mazdeh, M., Abolfathi, P., Mohammadi, Y., and Mehrpooya, M. (2020). Evidence for a Beneficial Effect of Oral N-acetylcysteine on Functional Outcomes and Inflammatory Biomarkers in Patients with Acute Ischemic Stroke. Neuropsychiatr Dis Treat 16, 1265–1278.

Salazar-Ramiro, A., Ramirez-Ortega, D., Perez de la Cruz, V., Hernandez-Pedro, N.Y., Gonzalez-Esquivel, D.F., Sotelo, J., and Pineda, B. (2016). Role of Redox Status in Development of Glioblastoma. Front Immunol 7, 156.

Satpathy, S., Krug, K., Jean Beltran, P.M., Savage, S.R., Petralia, F., Kumar-Sinha, C., Dou, Y., Reva, B., Kane, M.H., Avanessian, S.C., et al. (2021). A proteogenomic portrait of lung squamous cell carcinoma. Cell 184, 4348–4371.e4340.

Shin, C.S., Mishra, P., Watrous, J.D., Carelli, V., D’Aurelio, M., Jain, M., and Chan, D.C. (2017). The glutamate/cystine xCT antiporter antagonizes glutamine metabolism and reduces nutrient flexibility. Nat Commun 8, 15074.

Silver, D.J., Roversi, G.A., Bithi, N., Wang, S.Z., Troike, K.M., Neumann, C.K., Ahuja, G.K., Reizes, O., Brown, J.M., Hine, C., et al. (2021). Severe consequences of a high-lipid diet include hydrogen sulfide dysfunction and enhanced aggression in glioblastoma. J Clin Invest.

Singh, F., Charles, A.L., Schlagowski, A.I., Bouitbir, J., Bonifacio, A., Piquard, F., Krahenbuhl, S., Geny, B., and Zoll, J. (2015). Reductive stress impairs myoblasts mitochondrial function and triggers mitochondrial hormesis. Biochim Biophys Acta 1853, 1574–1585.

Smilkstein, M.J., Knapp, G.L., Kulig, K.W., and Rumack, B.H. (1988). Efficacy of oral N-acetylcysteine in the treatment of acetaminophen overdose. Analysis of the national multicenter study (1976 to 1985). N Engl J Med 319, 1557–1562.

Stupp, R., Mason, W.P., van den Bent, M.J., Weller, M., Fisher, B., Taphoorn, M.J., Belanger, K., Brandes, A.A., Marosi, C., Bogdahn, U., et al. (2005). Radiotherapy plus concomitant and adjuvant temozolomide for glioblastoma. N Engl J Med 352, 987–996.

Stupp, R., Taillibert, S., Kanner, A.A., Kesari, S., Steinberg, D.M., Toms, S.A., Taylor, L.P., Lieberman, F., Silvani, A., Fink, K.L., et al. (2015). Maintenance Therapy With Tumor-Treating Fields Plus Temozolomide vs Temozolomide Alone for Glioblastoma: A Randomized Clinical Trial. JAMA 314, 2535–2543.

Subramanian, A., Tamayo, P., Mootha, V.K., Mukherjee, S., Ebert, B.L., Gillette, M.A., Paulovich, A., Pomeroy, S.L., Golub, T.R., Lander, E.S., et al. (2005). Gene set enrichment analysis: a knowledge-based approach for interpreting genome-wide expression profiles. Proc Natl Acad Sci U S A 102, 15545–15550.

Tokumaru, S., Iguchi, H., and Kojo, S. (1996). Change of the lipid hydroperoxide level in mouse organs on ageing. Mech Ageing Dev 86, 67–74.

Tsai, J.C., Jain, M., Hsieh, C.M., Lee, W.S., Yoshizumi, M., Patterson, C., Perrella, M.A., Cooke, C., Wang, H., Haber, E., et al. (1996). Induction of apoptosis by pyrrolidinedithiocarbamate and N-acetylcysteine in vascular smooth muscle cells. J Biol Chem 271, 3667–3670.

Venable, J.H., and Coggeshall, R. (1965). A Simplified Lead Citrate Stain for Use in Electronmicroscopy. Journal of Cell Biology 25, 407–408.

Venkataramani, V., Tanev, D.I., Strahle, C., Studier-Fischer, A., Fankhauser, L., Kessler, T., Korber, C., Kardorff, M., Ratliff, M., Xie, R., et al. (2019). Glutamatergic synaptic input to glioma cells drives brain tumour progression. Nature 573, 532–538.

Venkatesh, H.S., Morishita, W., Geraghty, A.C., Silverbush, D., Gillespie, S.M., Arzt, M., Tam, L.T., Espenel, C., Ponnuswami, A., Ni, L., et al. (2019). Electrical and synaptic integration of glioma into neural circuits. Nature 573, 539–545.

Wang, L.-B., Karpova, A., Gritsenko, M.A., Kyle, J.E., Cao, S., Li, Y., Rykunov, D., Colaprico, A., Rothstein, J.H., Hong, R., et al. (2021). Proteogenomic and metabolomic characterization of human glioblastoma. Cancer Cell 39, 509–528.e520.

Weerapana, E., Wang, C., Simon, G.M., Richter, F., Khare, S., Dillon, M.B., Bachovchin, D.A., Mowen, K., Baker, D., and Cravatt, B.F. (2010). Quantitative reactivity profiling predicts functional cysteines in proteomes. Nature 468, 790–795.

Wickham, H., François, R., Henry, L., and Müller, K. (2021). dplyr: A Grammar of Data Manipulation.

Wrobel, M., Czubak, J., Bronowicka-Adamska, P., Jurkowska, H., Adamek, D., and Papla, B. (2014). Is development of high-grade gliomas sulfur-dependent? Molecules 19, 21350–21362.

Xu, R., Tao, A., Bai, Y., Deng, Y., and Chen, G. (2016). Effectiveness of N-Acetylcysteine for the Prevention of Contrast-Induced Nephropathy: A Systematic Review and Meta-Analysis of Randomized Controlled Trials. J Am Heart Assoc 5.

Yang, G., Zhao, K., Ju, Y., Mani, S., Cao, Q., Puukila, S., Khaper, N., Wu, L., and Wang, R. (2013). Hydrogen sulfide protects against cellular senescence via S-sulfhydration of Keap1 and activation of Nrf2. Antioxid Redox Signal 18, 1906–1919.

Yang, W., Zheng, Y., Xia, Y., Ji, H., Chen, X., Guo, F., Lyssiotis, C.A., Aldape, K., Cantley, L.C., and Lu, Z. (2012). ERK1/2-dependent phosphorylation and nuclear translocation of PKM2 promotes the Warburg effect. Nat Cell Biol 14, 1295–1304.

Ye, Z.C., and Sontheimer, H. (1999). Glioma cells release excitotoxic concentrations of glutamate. Cancer Res 59, 4383–4391.

Yu, Y., Villanueva-Meyer, J., Grimmer, M.R., Hilz, S., Solomon, D.A., Choi, S., Wahl, M., Mazor, T., Hong, C., Shai, A., et al. (2021). Temozolomide-induced hypermutation is associated with distant recurrence and reduced survival after high-grade transformation of low-grade IDH-mutant gliomas. Neuro Oncol.

Zhang, H., Limphong, P., Pieper, J., Liu, Q., Rodesch, C.K., Christians, E., and Benjamin, I.J. (2012). Glutathione-dependent reductive stress triggers mitochondrial oxidation and cytotoxicity. FASEB J 26, 1442–1451.

Zhao, L., Wang, Y., Yan, Q., Lv, W., Zhang, Y., and He, S. (2015). Exogenous hydrogen sulfide exhibits anti-cancer effects though p38 MAPK signaling pathway in C6 glioma cells. Biol Chem 396, 1247–1253.

Zhu, J., and Thompson, C.B. (2019). Metabolic regulation of cell growth and proliferation. Nat Rev Mol Cell Biol 20, 436–450.

